# Caveolin-1 Regulates Neurogenesis and Learning and Memory by Modulation of Mitochondrial Dynamics

**DOI:** 10.1101/2023.09.23.558792

**Authors:** Terilyn K. L. Stephen, Deepika Patel, Luis Aponte Cofresi, Elvis Quiroz, Kofi Owusu-Ansah, Yomna Ibrahim, Ellis Qualls, Jeffery Marshall, Wenping Li, Pavan Kumar, Aashutosh Shetti, Jacqueline A Bonds, Richard D. Minshall, Stephanie M. Cologna, Orly Lazarov

**Affiliations:** Department of Anatomy and Cell Biology, University of Illinois at Chicago, Chicago, IL, USA; Department of Chemistry, University of Illinois at Chicago, IL, USA; Departmet of Anesthesiology, University of California San Diego, CA, USA; Deparment of Pharmacology and Regenerative Medicine, University of Illinois at Chicago, IL, USA; Department of Anesthesiology, University of Illinois at Chicago, IL USA

**Keywords:** Caveolin-1, Adult Hippocampal Neurogenesis, Neural Stem Cells, Neuronal Differentiation, Mitochondria

## Abstract

Hippocampal neurogenesis plays instrumental roles in learning and memory. However, the mechanisms underlying neurogenesis are not fully understood. Here we show that the expression of Caveolin-1 (Cav-1), the principal component of caveolae, peaks in neural progenitor cells (NPCs) during neurogenesis. Using NestinCreER^T2^;Cav-1^fl/fl^ male mice, and CRISPR-sham (Cav-1 Ctrl) and CRISPR/Cas9-edited (Cav-1 KO) human induced-pluripotent stem cells, we observed that Cav-1 deletion led to reduced stem cell proliferation and enhanced differentiation into neurons. This was manifested by increased neuronal dendritic tree surface area and enhanced mouse performance in contextual discrimination. Proteomic analysis revealed that Cav-1 plays a role in mitochondrial pathways in NPCs. Cav-1 localized to the mitochondria in NPCs and co-immunoprecipitated with mitofusion 2. Mitochondrial morphology was elongated in Cav-1 KO NPCs and the expression of mitofusion 2 was increased in mitochondrial fractions. Restoration of Cav-1 levels rescued elongated mitochondrial morphology and altered neuronal differentiation in Cav-1 KO NPCs. Together, this study identifies Cav-1 as a novel regulator of neurogenesis and -dependent learning and memory.

**Significance Statement:** The hippocampal dentate gyrus (DG) orchestrates adult hippocampal neurogenesis (AHN). The precise mechanisms governing AHN remain elusive. Caveolin-1 regulates neurogenesis through mitochondrial fission-fusion process, suggesting Caveolin-1 as a novel regulator of AHN and underscoring the impact of AHN on cognition.

## Introduction

Adult hippocampus neurogenesis (AHN) is the postnatal continuous formation of new neurons and their functional integration in the granular cell layer of the dentate gyrus (DG) of the hippocampus. Radial glia-like neural stem cells (NSC) and neural progenitor cells (NPCs, together referred to NSPC) that reside in the subgranular zone (SGZ) of the DG, undergo proliferation and differentiation to produce new granule neurons (Toda et al., 2019). New neurons contribute to spatial and contextual learning and memory (Sahay et al., 2011; Niibori et al., 2012; Miller and Sahay, 2019; Masachs et al., 2021). However, AHN declines with age. In mice, marked reductions in NSPC proliferation and neurogenesis are observed as early as 6 months of age (Encinas et al., 2011; Demars et al., 2013; Britton et al., 2022). Impaired AHN occurs in various brain disorders, including Alzheimer’s Disease (AD), both in animal models and humans (Demars et al., 2010; Moreno-Jimenez et al., 2019; Tobin et al., 2019; Terreros-Roncal et al., 2021). In human cohorts the number of neuroblasts is inversely correlated with cognitive diagnosis (Tobin et al., 2019). Interesting, strategies that enhance neurogenesis, such as genetic and pharmacologic interventions restore memory deficits in AD mouse models (Hu et al., 2010; Choi et al., 2018; Mishra et al., 2022). While many intrinsic and extrinsic factors modulate AHN (Demars et al., 2011; Gadadhar et al., 2011; Artegiani et al., 2017; Vicidomini et al., 2020; Zhou et al., 2022), the mechanisms underlying the maintenance of NSPC and neurogenesis-dependent memory function are still not fully understood.

Caveolin-1 (Cav-1) is a 21-24 kDa scaffolding protein of the caveolin gene family that is crucial for the formation of caveolae (Williams and Lisanti, 2004). Cav-1 governs numerous signaling pathways (Parton, 2018; Parton et al., 2020). Cav-1 is enriched in endothelial cells (Ikezu et al., 1998; Virgintino et al., 2002) and is essential for blood brain barrier integrity and neurovascular coupling (Knowland et al., 2014; Blochet et al., 2020; Chow et al., 2020; Filchenko et al., 2020).

Previously, we showed that Cav-1 expression is depleted in the hippocampus of a Type II diabetes mouse model, and restoration of Cav-1 improved hippocampal memory. Cav-1 knockout mice show diverse neurological deficits, including impaired cholinergic function and hippocampal plasticity (Trushina et al., 2006; Gioiosa et al., 2008; Head et al., 2010). Overexpressing Cav-1 in hippocampal neurons enhances dendritic growth and arborization, likely via increased lipid raft formation and enhanced trafficking of synaptic receptors (Head et al., 2011; Egawa et al., 2018). Few studies exist examining Cav-1 in neurogenic cell types. In the developing cortex, Cav-1 expression is essential for internalization of cell adhesion proteins that regulate proper migration and dendritic pruning in immature neurons (Shikanai et al., 2018). In early stages of neuronal differentiation from NPCs derived from human inducible pluripotent stem cells (iPSCs), Cav-1 phosphorylation at its tyrosine 14 site is needed for axonal growth (Wang et al., 2019). However, a role for Cav-1 in AHN has never been described. Here, we show that Cav-1 is expressed in both hippocampal NSPC isolated from mice and human NPCs and neurons derived from iPSCs. Cell-specific deletion of Cav-1 in AHN in mice, resulted in significant reductions in NSC proliferation, increased neuronal differentiation and enhanced performance of mice in the AHN-dependent contextual discrimination. Similarly, Cav-1 deletion from iPSCs induced NPCs resulted in stalled proliferation and enhanced mature neuron formation. Proteomic analysis revealed that Cav-1 regulates AHN via mitochondria-related protein pathways. Prior reports linked mitochondrial fusion to neuronal differentiation (Khacho and Slack, 2018; Kochan et al., 2024). However, the molecular signals that regulate this process is largely unknown. Importantly, we observed that Cav-1 localizes to the mitochondria in iPSCs-derived NPCs and regulates fission fusion via interaction with mitofusion-2. Reinstating Cav-1 in Cav-1 KO iPSCs-derived NPCs restored structural alterations in mitochondria. This study determines that Cav-1 is a novel regulator of AHN in the mouse and human brain and governs neuronal differentiation.

## Results

### Cav-1 deletion in NSPCs shift neurogenesis towards neuronal differentiation in mice

Nestin expression is high in actively proliferating NSPC and downregulated as cells exit the cell cycle during differentiation (Frederiksen and McKay, 1988; Wilhelmsson et al., 2019). To investigate the role of Cav-1 in AHN, we generated a NSPC specific inducible Cav-1 knockout model, where we bred a Cav-1 floxed mice (Cav-1^fl/fl^) (Cao et al., 2003; Oliveira et al., 2017) with a tamoxifen inducible Cre recombinase driven by a Nestin promoter (NestinCre^ERT2/+^) (Mishra et al., 2022). NestinCre^ERT2/+^;Cav-1^fl/fl^ mice were injected at 4-5 weeks of age with either tamoxifen to conditionally delete Cav-1 from NSPCs (iNSC Cav-1 KO) or with corn oil to generate control mice (iNSC Cav-1 WT; **Figure S1 A)**. We confirmed Cav-1 recombination in hippocampal NSPC isolated from iNSC Cav-1 KO and iNSC Cav-1 WT mice by quantitative real-time PCR (qPCR), immunoblotting and immunostaining **(Figure S1 B-D).** Ultrastructural transmission electron microscopy was used to determine if Cav-1 expression in NSPC correlated with the presence of caveolae. Flask-shaped invaginations and vesicles resembling caveolae were identified in the iNSC Cav-1 WT NSPC, whereas larger (>150 nm) electron dense vesicles resembling clathrin-coated vesicles were predominantly found in the iNSC Cav-1 KO NSPC **(Figure S1 E).** Together, this indicates that Cav-1 is expressed in adult hippocampal NSPCs, associated with the presence of caveolae, and that tamoxifen-induced recombination leads to a complete deletion of Cav-1 in iNSC Cav-1 KO mice.

To assess the impact of Cav-1 deletion on neurogenesis, we analyzed NSC proliferation in the DG of iNSC Cav-1 WT and KO mice. At both 3 and 6 months of age, there were no significant differences in the total NSC population (GFAP^+^Nestin^+^) or in the quiescent NSC subpopulation (GFAP^+^Nestin^+^MCM2-) between genotypes (**Figure 1 A-C and 1 E-G)**. However, the number of proliferating NSCs (GFAP^+^Nestin^+^MCM2^+^) was significantly reduced in Cav-1 KO mice at both ages (**Figure 1 D and 1 H)**. In line with this, expression of ASCL-1 and TBR-2-proxies of proliferating NSCs and intermediate progenitor cells (IPCs), respectively-was also significantly reduced in Cav-1 KO mice, indicating impaired expansion of these cell populations at both ages (**Figure S1 F-M).** Supporting these in vivo findings, neurosphere assays revealed that Cav-1 KO NSPCs had reduced clone formation, smaller clone size, fewer total cells (**Figure 1 I-L**) and reduced incorporation of EDU compared to WT controls **(Figure S1 N-O).**

**Figure 1.**
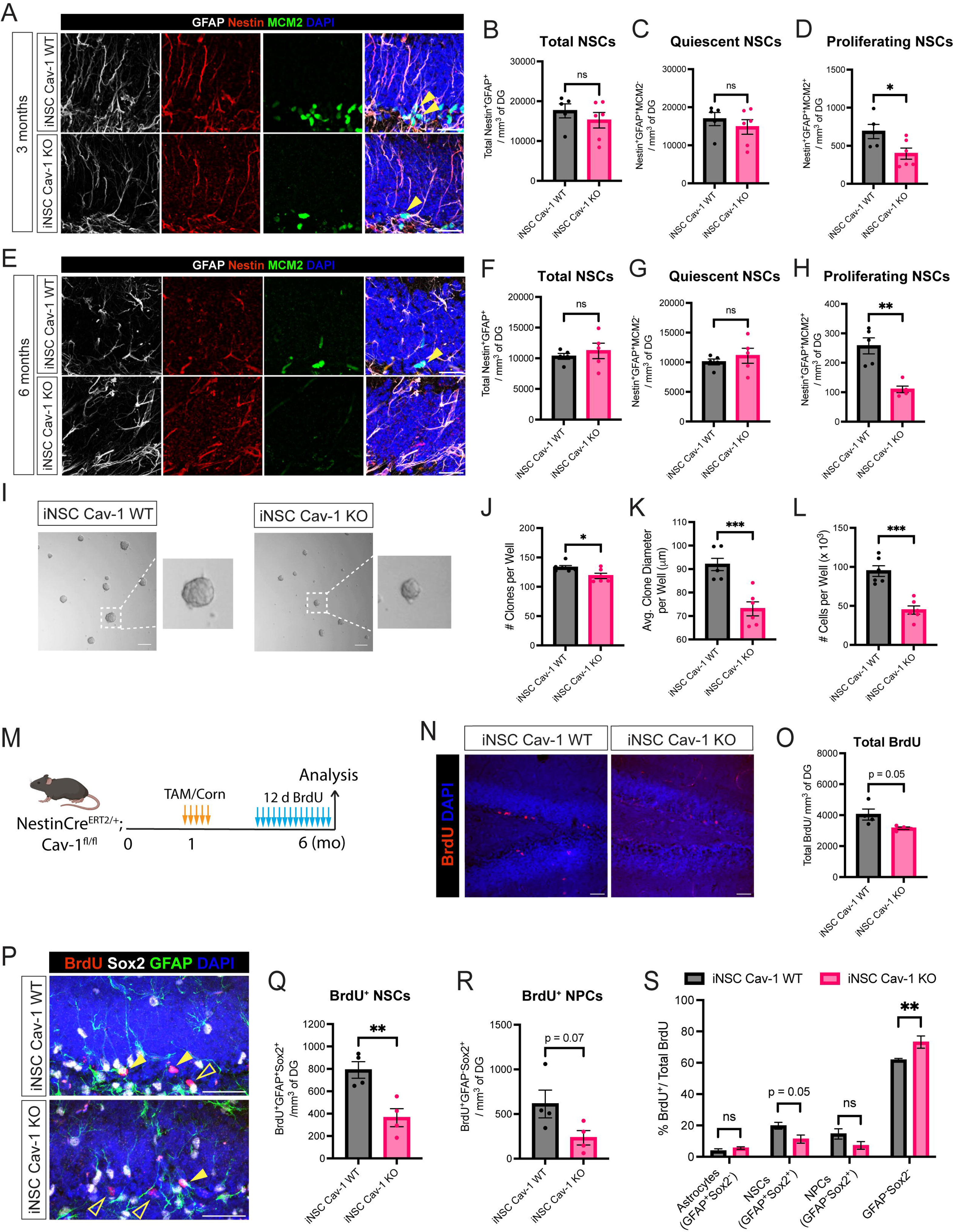
iNSC Cav-1 KO mice have reduced levels of proliferating NSPC in the DG. (A) Representative confocal images of GFAP (white), Nestin (red), MCM2 (green) and DAPI (blue) markers in the DG of iNSC Cav-1 WT and iNSC Cav-1 KO mice at 3 months of age. Yellow arrowheads indicate GFAP^+^Nestin^+^MCM2^+^ cells. Scale bar, 25 μm. (B-D) Quantification of total NSC (GFAP^+^Nestin^+^), quiescent NSC (GFAP^+^Nestin^+^MCM2^-^) and proliferating NSC (GFAP^+^Nestin^+^MCM2^+^) in the DG of iNSC Cav-1 WT and iNSC Cav-1 KO mice at 3 months of age. n=5 mice per group. (E) Representative confocal images of GFAP (white), Nestin (red), MCM2 (green) and DAPI (blue) positive cells in the DG of iNSC Cav-1 WT and iNSC Cav-1 KO mice at 6 months of age. Yellow arrowheads indicate GFAP^+^Nestin^+^MCM2^+^ cells. Scale bar, 25 μm. (F-H) Quantification of total NSC (GFAP^+^Nestin^+^), quiescent NSC (GFAP^+^Nestin^+^MCM2^-^) and proliferating NSC (GFAP^+^Nestin^+^MCM2^+^) in the DG of iNSC Cav-1 WT and iNSC Cav-1 KO mice at 6 months of age. n=5 mice per group. I-L) Clonogenic proliferation assay in NSPC isolated neurospheres from iNSC Cav-1 WT and iNSC Cav-1 KO mice. The number of clones (neurospheres), average clone diameter and number of cells after dissociation of clones on day 5 of the assay were quantified per well. Scale bar, 100 μm. (M) 5-bromo-2’-deoxyuridine (BrdU) pulse strategy to quantify changes in NSC fate and differentiation in DG of iNSC Cav-1 WT and iNSC Cav-1 KO mice at 6 months of age. Mice were injected daily for 12 consecutive days and sacrificed 24 hr after the last injection. (N-O) Confocal images of BrdU immunostaining and quantification total number of BrdU^+^ cells in the DG of 6-month-old iNSC Cav-1 KO and iNSC Cav-1 WT mice. Scale bar, 25 μm. n=4 mice per group. (P) Representative confocal images of BrdU (red), Sox2 (white), GFAP (green) and DAPI (blue) markers in the DG of iNSC Cav-1 WT and iNSC Cav-1 KO mice at 6 months of age. Solid yellow arrowheads indicate BrdU^+^GFAP^+^Sox2^+^ cells. Outlined yellow arrowheads indicate BrdU^+^GFAP^-^Sox2^-^ cells. Scale bar, 25 μm. (Q-S) Quantification of total BrdU^+^ NSCs (GFAP^+^Sox2^+^), BrdU^+^ NPCs (GFAP^+^Sox2^-^) and percentage of BrdU^+^ cell phenotypes normalized to total number of BrdU ^+^ cells in the DG of iNSC Cav-1 WT and iNSC Cav-1 KO mice. n=4 mice per group. Data represented as mean ± SEM. Data analyzed by unpaired two-tailed Student’s t-test except (S) which was analyzed by two-way ANOVA with Tukey multiple comparisons correction. ns p> 0.05, *p< 0.05, **p < 0.01, ***p < 0.001.

We next utilized a 12 day 5-bromo-2’-deoxyuridine (BrdU) pulse labeling paradigm followed by analysis 24 hr after the final BrdU injection to assess NSPC proliferation and lineage progression (**Figure 1 M**). A trending reduction (p = 0.05) in the total level of BrdU positive cells in the DG was observed in the iNSC Cav-1 KO mice compared to the iNSC Cav-1 WT mice (**Figure 1 N-O**). Consistent with our proliferation analysis, BrdU⁺ GFAP⁺ Sox2⁺ NSCs were significantly reduced in iNSC Cav-1 KO mice (**Figure 1 P-Q**), and a trend toward fewer BrdU⁺GFAP⁻Sox2⁺ NPCs was also detected **(p = 0.07; Figure 1 R)**. More notably, there was a significant shift in the lineage of BrdU^+^ cells away from NSPCs pool in the iNSC Cav-1 KO mice, suggesting that Cav-1 loss alters early neurogenic fate decisions (**Figure 1 S)**.

Given that stalled proliferation of NSC can influence early stages of differentiation, we quantified triple-labeled BrdU⁺DCX⁺NeuN⁻ neuroblasts and BrdU⁺DCX⁺NeuN⁺ immature neurons (**Figure 2 A-D**). While no significant differences were observed in BrdU⁺ neuroblasts (**Figure 2 B)**, the number of BrdU⁺ immature neurons were significantly elevated in Cav-1 KO mice compared with WT (**Figure 2 C-D**). Consistent with these findings, at 6 months of age the iNSC Cav-1 KO mice exhibited a significant increase in total number of DCX^+^ NPCs, neuroblasts and immature neurons (DCX^+^NeuN^+^) compared to iNSC Cav-1 WT mice (**Figure 2 E-H, Figure S1 P-S).** These findings support the hypothesis that Cav-1 loss promotes neuronal lineage commitment.

**Figure 2.**
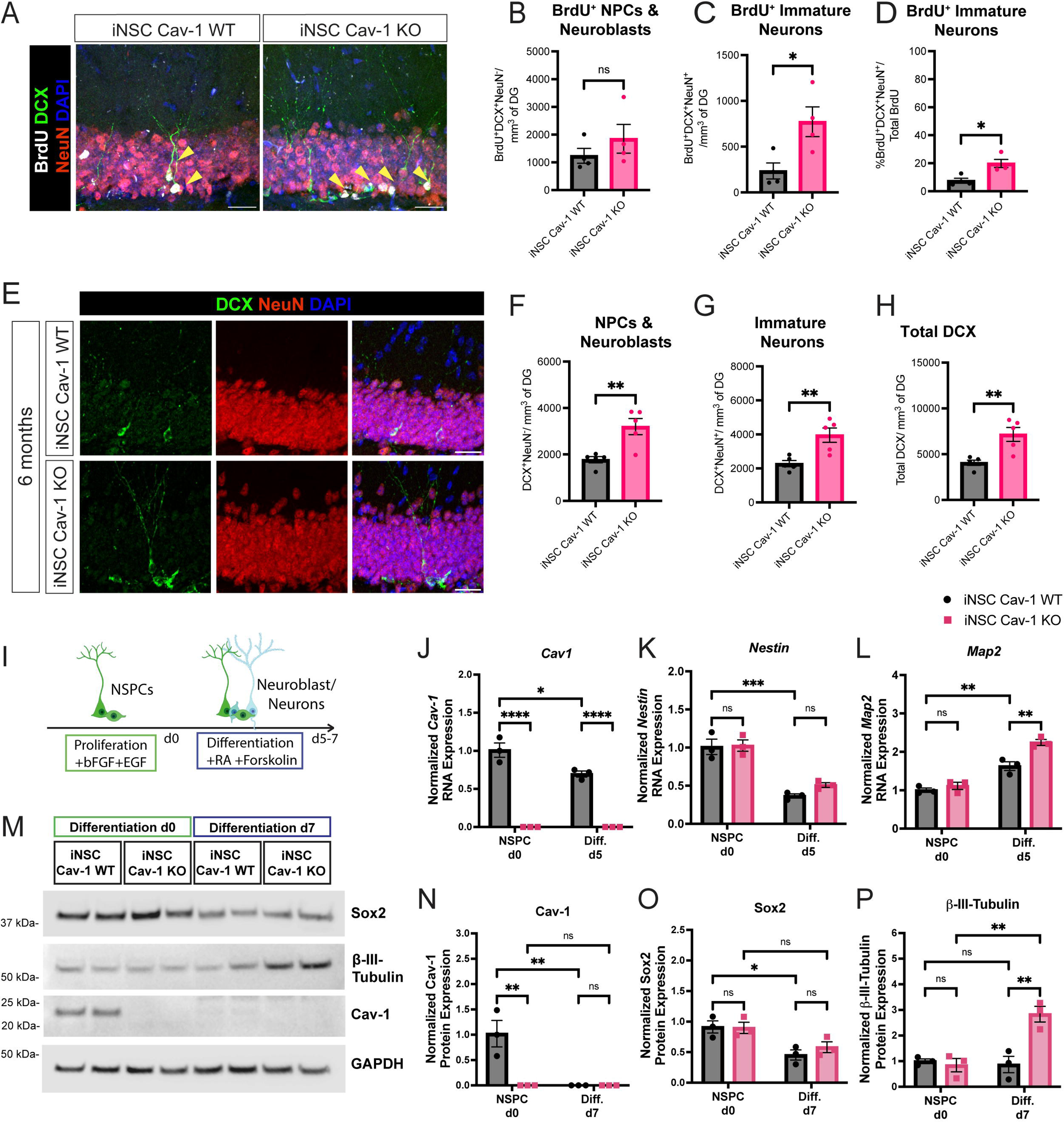
Cav-1 regulates *in vivo* and *in vitro* differentiation of hippocampal NSPCs (A) Representative confocal images of BrdU (white), DCX (Green), NeuN (red) and DAPI (blue) markers in the DG of iNSC Cav-1 WT and iNSC Cav-1 KO mice at 6 months of age. Outlined yellow arrowheads indicate BrdU^+^DCX^+^NeuN^+^ cells. Scale bar, 25 μm. (B-D) Quantification of BrdU^+^ DCX expressing NPCs and neuroblasts (DCX^+^NeuN^-^), immature neurons (DCX^+^NeuN^+^), and percentage of BrdU^+^ immature neurons (DCX^+^NeuN^+^) normalized to total number of BrdU^+^ cells in the DG of iNSC Cav-1 WT and iNSC Cav-1 KO mice. n=4 mice per group. (E) Confocal images of DCX and NeuN immunostaining in the DG of iNSC Cav-1 WT and iNSC Cav-1 KO mice at 6 months of age. Scale bar, 25 μm. (F-H) Quantification of NPCs and neuroblasts (DCX^+^NeuN^-^) and immature neurons (DCX^+^NeuN^+^) and total DCX expressing cells in the DG of iNSC Cav-1 WT and iNSC Cav-1 KO mice. n=4 mice per group. (I) Schematic representation of the protocol used for neural differentiation of primary hippocampal NSPCs. (J-L) Quantification of Cav-1, Nestin, and MAP2 transcript expression by RT-qPCR between iNSC Cav-1 WT and iNSC Cav-1 KO NSPCs undergoing differentiation for 5 days. Expression level normalized to d0 of iNSC Cav-1 WT NSPC (n=3 replicates). (M) Immunoblot of Cav-1, Sox2 and β-III-Tubulin in NSPC isolated from iNSC Cav-1 KO and iNSC Cav-1 WT mice undergoing differentiation for 7 days. (N-P) Quantification of Cav-1, Sox2 and β-III-Tubulin normalized to GAPDH expression and then normalized to d0 iNSC Cav-1 WT NSPC (n=3 replicates). Data represented as mean ± SEM. Data analyzed by unpaired two-tailed Student’s t-test (B-D and F-H) and two-way ANOVA with Tukey multiple comparisons correction (J-L and N-P). ns p > 0.05, *p < 0.05, **p < 0.01, ***p < 0.001, and ****p < 0.0001.

The effect of Cav-1 deletion on differentiation was also examined in hippocampal neurosphere cultures isolated from iNSC Cav-1 WT and iNSC Cav-1 KO mice (**Figure 2 I**) (Bonds et al., 2020; Gadadhar et al., 2011). During in vitro differentiation over 5–7 days, Cav-1 transcript and protein levels were significantly downregulated in WT NSPCs (**Figure 2 J, M-N).**

Differentiation progression was confirmed by decreased expression of nestin and Sox2 and concurrent upregulation of the neuronal markers Map2 and βIII-tubulin (**Figure 2 K-L, O-P**). Consistent with in vivo findings, Cav-1 KO NSPCs exhibited significantly higher expression of Map2 and βIII-tubulin compared with WT controls during differentiation (**Figure 2 L, P).** Together these findings suggest that Cav1 negatively regulates NSPC differentiation. Its loss facilitates the transition from proliferation to neuronal commitment, enhancing the maturation of new neurons.

### Cav-1 deletion in NSPC increases newborn neuron dendritic tree volume and improves context discrimination learning and memory

Dendritic arborization of immature neurons is a critical component of neuronal maturation, influencing the ability of these adult-born neurons to integrate into existing circuits and participate in hippocampal-dependent functions such as learning and memory (Toni et al., 2007; Zhao et al., 2008). To investigate whether Cav-1 deletion alters dendritic structure in newly generated neurons, we infected dentate gyrus neurons in iNSC Cav-1 WT and iNSC Cav-1 KO mice with retrovirus-green fluorescence protein (RV-GFP) and sacrificed them 30 days later to allow the visualization of the dendritic structure of newly post-mitotic neurons. Dendrite morphology was quantified by Sholl analysis **(Figure 3A).** We observed no significant differences in the number of dendritic branch intersections and total dendritic length (**Figure 3 B-D**). Yet, both surface area and more significantly, dendritic volume were increased in the RV-GFP labeled newly born neurons in the iNSC Cav-1 KO mice compared to iNSC Cav-1 WT (**Figure 3 E-F**). These findings suggest that Cav-1-deficient newborn neurons, despite unchanged dendritic branching patterns, develop thicker dendrites with potentially greater membrane and cytoplasmic volume. These features are often associated with enhanced structural stability and increased potential for synaptic integration (van Elburg and van Ooyen, 2010).

**Figure 3.**
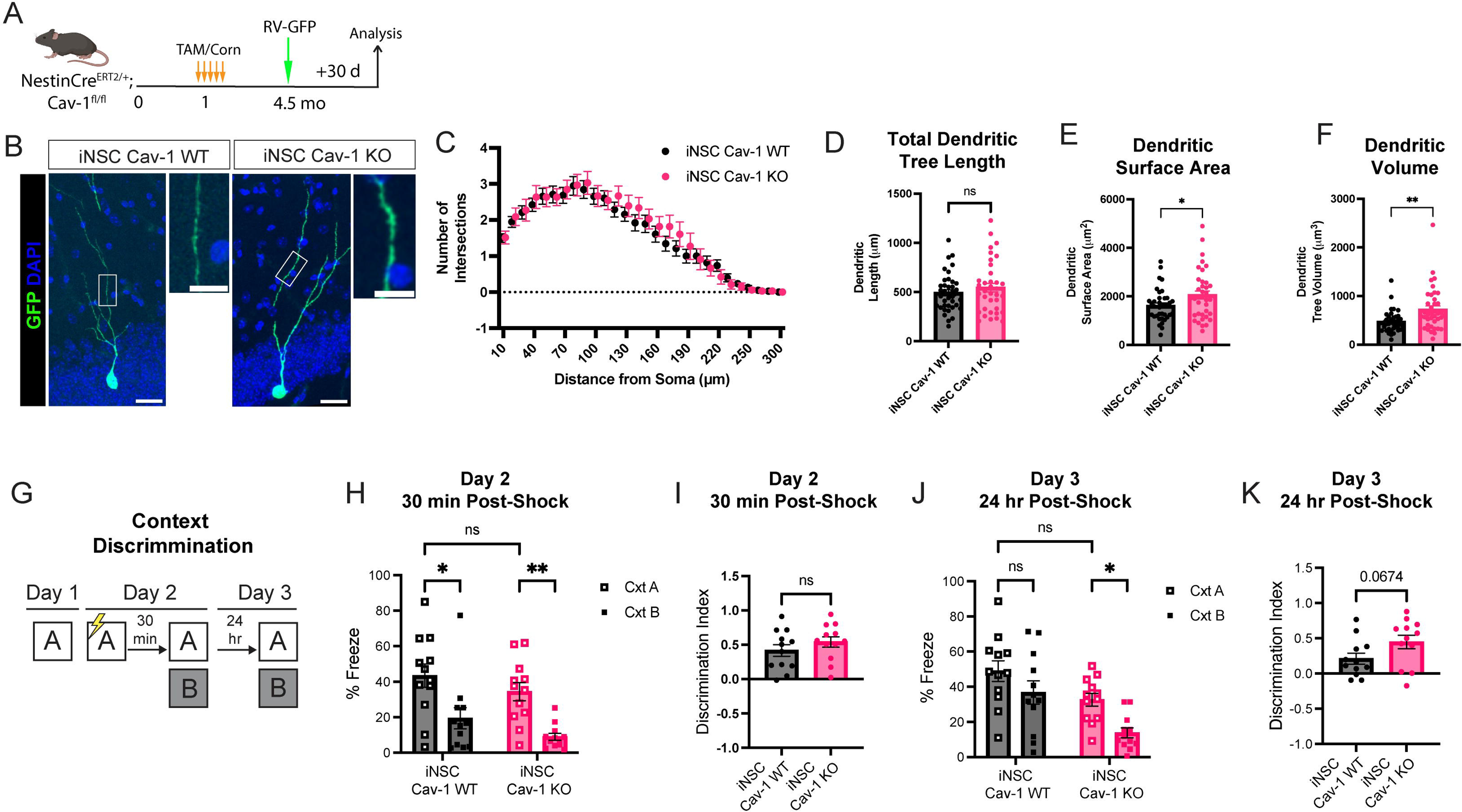
iNSC Cav-1 KO mice display enhanced dendritic volume of newborn neurons and improved contextual discrimination learning and memory. (A) GFP retrovirus (RV-GFP) labeling strategy of newborn neurons. Mice were stereotaxic injected with RV-GFP in the DG and then, sacrificed 30 days post injection for analysis. (B) Confocal images of GFP^+^ newborn neurons with zoomed dendritic tree arborization in the DG of 5.5-month-old iNSC Cav-1 KO and iNSC Cav-1 WT mice. Scale bar, 25 μm and 10 μm. (C) Sholl analysis data represented as mean number of intersections for each successive 30Lμm segment as a function of the radial distance of the corresponding segment from the soma. (D-F) Quantification of total dendritic tree length, surface area and volume respectively. n=4 mice per group, 28-30 cells analyzed total per group. (G) Schematic of contextual fear discrimination paradigm between Context A and B. See Materials and methods for details. (H-I) Quantification of percent freeze percent freeze (30 min post-shock) and discrimination index in context A and B on Day 2 between the iNSC Cav-1 WT and iNSC Cav-1 KO mice. n=12 per genotype. (J-K) Quantification of percent freeze (24 hr post-shock) and discrimination index in context A and B on Day 3 between the iNSC Cav-1 WT and iNSC Cav-1 KO mice. n=12 per genotype. Data represented as mean ± SEM. Data analyzed in (C) by mixed model a mixed effects model with the Geisser-Greenhouse correction followed by Bonferroni post-hoc test for multiple comparisons; in D-F, I and K analyzed by unpaired two-tailed Student’s t-test; and in H and J by a two-way ANOVA with Tukey’s multiple comparisons correction. ns p > 0.05, *p < 0.05, **p < 0.01.

Given that newborn neurons (4-6 weeks post-mitotic) become selectively activated during memory formation and are essential for contextual discrimination, the ability to discriminate between similar contexts (Clelland et al., 2009; Sahay et al., 2011; Nakashiba et al., 2012), we next asked whether the increase in immature neurons in the iNSC Cav-1 KO mice results in improved neurogenesis-dependent learning and memory. Thus, 6 month old iNSC Cav-1 KO and iNSC Cav-1 WT mice underwent a contextual discrimination test (Clemenson et al., 2014) (**Figure 3 G).** Both iNSC Cav-1 KO and iNSC Cav-1 WT mice displayed higher freezing levels in Context A (Cxt A) compared to Context B (Cxt B) at 30 mins post-shock, albeit with no difference in the discrimination index (**Figure 3 H-I**). Interestingly 24 hours post-shock on Day 3, the iNSC Cav-1 KO mice had a significantly higher level of freezing in Cxt A compared to Cxt B whereas the iNSC Cav-1 WT mice exhibited the equal amount of freezing in both Cxt A and Cxt B (**Figure 3 J)**. A trending increase (p = 0.06) in the discrimination index was observed in the iNSC Cav-1 KO mice (**Figure 3 K**), suggesting enhanced neurogenesis-dependent memory.

We then examined whether the iNSC Cav-1 KO mice displayed enhanced context generalization compared to the iNSC Cav-1 WT mice 24 hours post-shock. Studies show that significant alterations in context geometry are sufficient to induce generalization rather than discrimination behavior (Gonzalez et al., 2003; Huckleberry et al., 2016) and that newborn neurons maintain generalization behavior (Guo et al., 2018). To evaluate context generalization, a similar paradigm was used with a Context C (Cxt C), which is modified in geometry and tactile cues **(Figure S2 A)**. Notably on Day 2 (30 mins post shock), iNSC Cav-1 WT mice were unable to discriminate Cxt A from Cxt C whereas iNSC Cav-1 KO mice had a significantly higher level of freezing in Cxt A compared to Cxt C with a trending significant discrimination index **(Figure S2 B-C)**. Notably, 24 hours post shock, both groups of mice were able to equally discriminate Cxt A from Cxt C 24 hr post-shock **(Figure S2 D-E).** No differences in level of anxiety-like behavior **(Figure S2 F-I)** nor performance in the spatial novel object location (NOL) task (**Figure SS J-L**) were observed between the iNSC Cav-1 KO and iNSC Cav-1 WT mice. Together these findings indicate that Cav-1 deletion in NSPCs promotes the generation of immature neurons with enhanced dendritic volume, leading to improved neurogenesis-dependent context discrimination without affecting general anxiety or spatial memory.

### Proteomic analysis reveals that Cav-1 regulates mitochondrial signaling pathways and morphology in hippocampal NSPCs

Building on our findings that Cav-1 deletion in NSPCs enhances hippocampal neurogenesis, we hypothesized that Cav-1 regulates pathways involved cell cycle cessation and differentiation. Using quantitative proteomics, we assessed the molecular mechanisms by which Cav-1 regulates adult hippocampal neurogenesis in mice (**Figure 4 A).** For that, we isolated hippocampal NSPCs from the dentate gyrus of iNSC Cav-1 KO and iNSC Cav-1 WT mice and examined their protein profile by mass spectrometry. We identified 4,730 proteins, with 326 differentially expressed (DEPs; ANOVA, p < 0.05, **Figure 4 B Table S1**) — 228 downregulated and 98 upregulated in Cav-1 KO NSPCs. Ingenuity Pathway Analysis (IPA) and Gene Ontology (GO) revealed that the top altered pathways included Mitochondrial Dysfunction, TCA Cycle II, Cholesterol Biosynthesis, and Oxidative Phosphorylation (**Figure 4 C-D, Table S2**). GO analysis also pointed to changes in ATP biosynthesis, the electron transport chain (ETC), mitochondria and ribosome subunits, and lipid oxidation.

**Figure 4.**
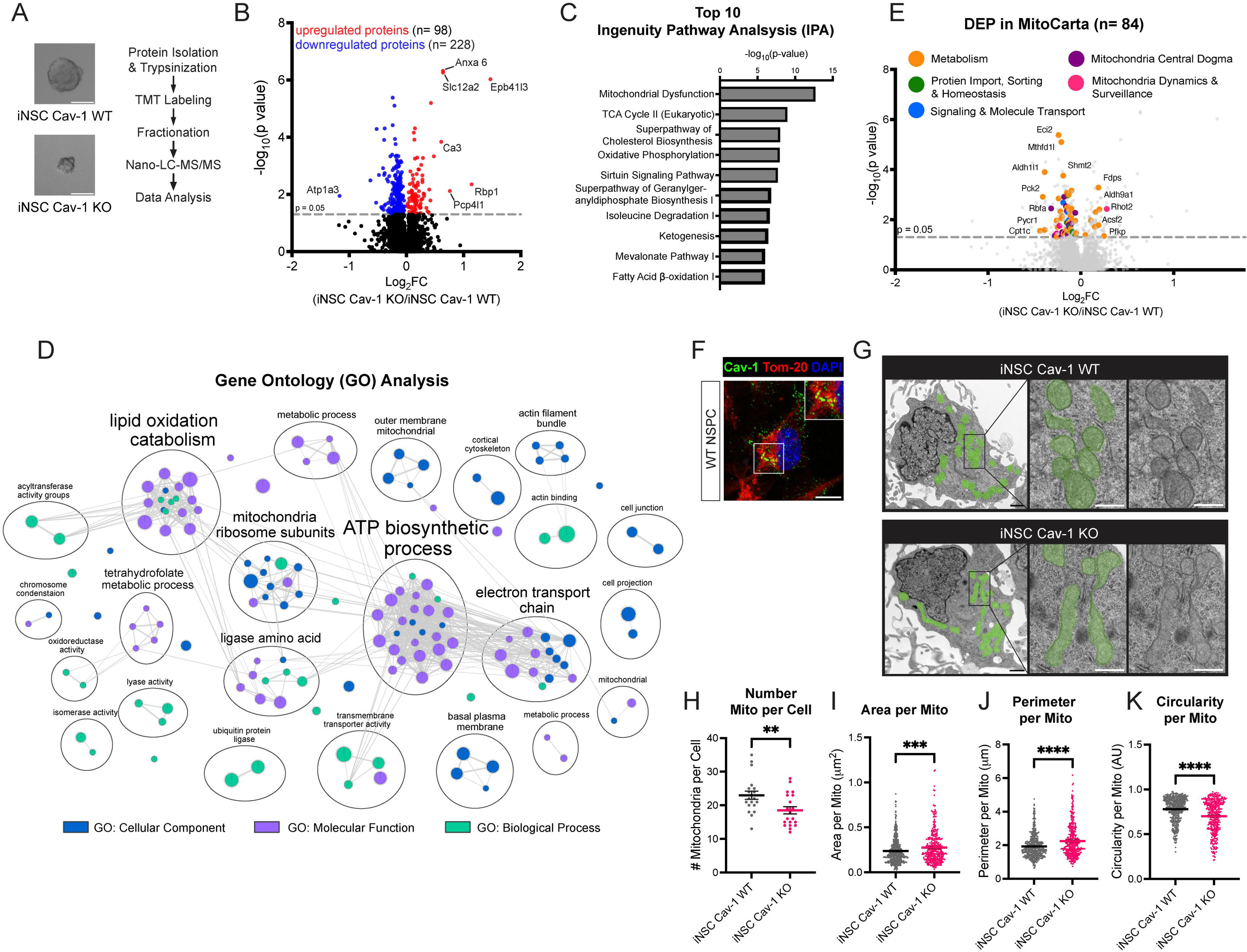
Loss of Cav-1 in mouse hippocampal NSPCs alters mitochondrial protein pathway expression and promotes mitochondrial elongation. (A) Schematic of proteomic workflow for hippocampal NSPCs isolated from iNSC Cav-1 KO and iNSC Cav-1 WT mice. See Materials and methods for details. (B) Volcano plot of proteins identified in iNSC Cav-1 KO vs iNSC Cav-1 WT cells. Significantly upregulated proteins in red and significantly downregulated proteins in blue. Data represented as Log_2_FoldChange (FC) of mean abundance of proteins normalized to iNSC Cav-1 WT (n=3 iNSC Cav-1 KO and n=3 iNSC Cav-1 WT). DEP determined by a one-way ANOVA with a p-value of < 0.05. (C) Bar chart showing the top 10 Ingenuity Pathway Analysis (IPA) altered pathways in the iNSC Cav-1 KO compared to iNSC Cav-1 WT hippocampal NSPCs (ANOVA, p < 0.05). (D) Cytoscape cluster mapping of altered GO pathways in hippocampal NSPCs isolated from iNSC Cav-1 KO and iNSC Cav-1 WT mice. Analysis based on Log_2_FC ratio of DEPs with a significant cut off value of p < 0.05. (E) Volcano plot of Mitocarta 3.0 (Rath et al., 2021) proteins identified in iNSC Cav-1 KO vs iNSC Cav-1 WT cells. Metabolism related protein in orange; Protein Import, Sorting & Homeostasis in green; Signaling and Molecule Transport in blue, Mitochondria Central Dogma in purple; and Mitochondria Dynamics and Surveillance in pink. (F) Representative confocal image of Caveolin-1 (Cav-1, Green), Mitochondrial import receptor subunit TOM20 homolog (Tom-20, green) and DAPI (blue) immunostaining in WT hippocampal NSPCs. Scale bar 10 μm. (G) Representative transmission electron microscopy (TEM) images of NSPCs isolated from iNSC Cav-1 KO and iNSC Cav-1 WT mice. Mitochondria are shaded in green; scale bar = 1 μm with zoomed image scale bar = 500 nm. (H-K) Quantification of the number of mitochondria (mito) per cell imaged, area per mitochondria, perimeter per mitochondria and circularity per mitochondria using ultrastructural electron microscopy. n=20 cells per genotype with n=460 mitochondria for iNSC Cav-1 WT and n=370 mitochondria for iNSC Cav-1 KO analyzed. Data represented as mean ± SEM. Data analyzed by unpaired two-tailed Student’s t-test. **p < 0.01, ***p < 0.001, and ****p < 0.0001.

To explore Cav-1’s role in mitochondrial regulation in NSPCs, we compared the DEPs to the MitoCarta3.0 gene dataset (Rath et al., 2021), which catalogs 1,140 mouse genes localized to mitochondria. Of the 326 DEPs identified in iNSC Cav-1 KO vs. WT NSPCs, 84 (24%) matched MitoCarta3.0 genes (**Figure 4 E**). Downregulated proteins in Cav-1 KO NSPCs were enriched in pathways related to mtDNA maintenance, mtRNA processing and translation, protein import, sorting, and mitochondrial homeostasis. Several metabolic pathways including glycolysis, TCA, OXPHOS, lipid, nucleotide, amino acid, vitamin, and metal metabolism were also downregulated (**Figure 4 E**). Noticeably, we found that mitochondrial dynamics and surveillance associated proteins including Metaxin-2 (MTX2) and Voltage-dependent Anion-selective Channel (Vdac1) were downregulated, whereas Mitochondrial Rho GTPase 2 (Rhot2, also called Miro2) was significantly enriched in the iNSC Cav-1 KO NSPCs compared to iNSC Cav-1 WT NSPCs (**Figure 4 E)**. Miro2 interacts with mitofusion-2 (Mfn-2) to coordinate fission/fusion events (Saotome et al., 2008) and in neurons, Miro2 is required for retrograde trafficking of mitochondria (Misko et al., 2010; Ruggiero et al., 2017) suggesting that part of Cav-1’s function may involve regulating mitochondrial positioning and dynamics in NPCs.

In hippocampal neurogenesis, changes in mitochondrial morphology and dynamics have been found to govern self-renewal and cell fate (Khacho et al., 2016). We first examined the co-localization of Cav-1 to mitochondria. Caveolae and caveolin-1 were previously shown to localize to the outer membrane of mitochondria (Patel and Insel, 2009; Fridolfsson et al., 2012). Cav-1 co-localizes with Mitochondrial import receptor subunit TOM20 homolog (Tom-20), a translocase located on the mitochondrial outer membrane mitochondria, in WT hippocampal NSPCs by utilizing 3D confocal microcopy (**Figure 4 F**). Next, we examined whether Cav-1 deletion in hippocampal NSPCs alters mitochondrial morphology by Ultrasound EM and live cell imaging analysis. Ultrastructural EM analysis revealed that Cav-1 KO NSPCs had fewer mitochondria per cell, yet significantly increased mitochondrial area and perimeter, with reduced circularity indicating a shift toward elongated morphology (**Figure 4 G-K).** Live cell imaging using TMRM confirmed these findings: Cav-1 KO mitochondria were larger (**Figure S3 A-C**), more elliptical (**Figure S3 D**), and more branched (**Figure S3 E**), with elevated TMRM fluorescence (**Figure S3 F**) indicating a hyperpolarized membrane potential. Time-lapse imaging of mitochondria was analyzed using the Trackmate plugin in ImageJ (Ershov et al., 2022) (**Figure S3 G**). Mitochondria in the iNSC Cav-1 KO NSPC had a significant increase in velocity and total distance traveled compared to iNSC Cav-1 WT NSPCs **(Figure S3 H-I**). Together, these data suggest that Cav-1 regulates mitochondrial structure and dynamics in hippocampal NSPCs. Its deletion promotes elongation, branching, and mobility, features of fusion and fission dynamics, potentially driving neuronal differentiation (Khacho and Slack, 2018).

### Cav-1 deletion enhances differentiation of human iPSCs-derived NSCs

To explore whether Cav-1 plays similar role in human neurogenesis, we next utilized iPSCs model. Using this approach, we examined the effects of Cav-1 deletion on neuronal lineage progression and maturation across defined stages of neural development via generation of a CRISPR/Cas9-edited Cav-1 knockout iPSCs (Cav-1 KO). Cav-1 Ctrl and Cav-1 KO iPSCs were differentiated into forebrain neurons using neural induction method (**Figure 5 A).** This method defines 3 neuronal differentiation stages, with neural progenitors (NPCs) having the most stemness, precursors cells resembling early stages of neuron lineage (similar to neuroblasts) and lastly neurons.

**Figure 5.**
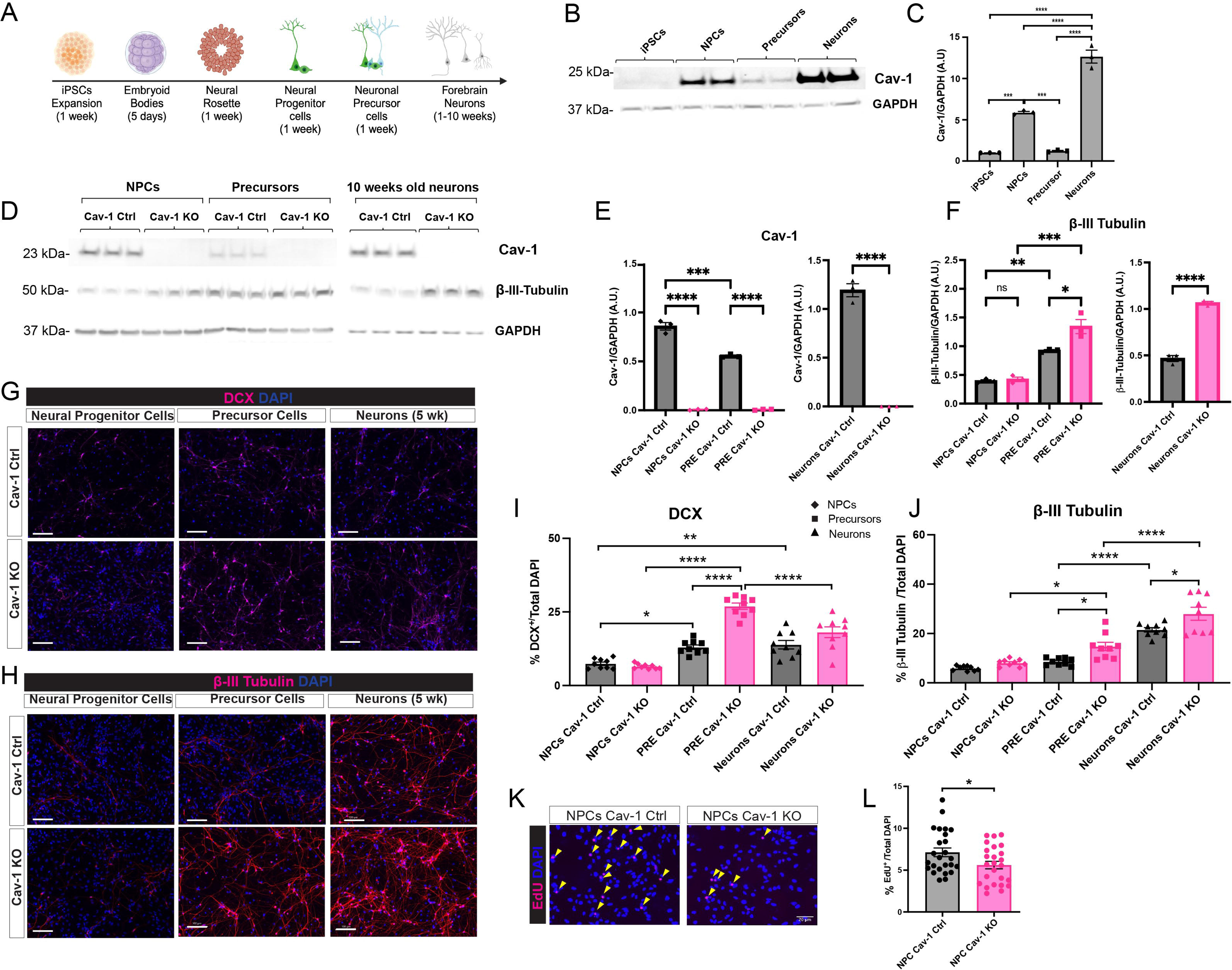
Cav-1 regulates proliferation and differentiation in human iPSCs-derived NPCs. (A) Schematic representation of the protocol used for neural induction of human induced-pluripotent stem cells (iPSCs) to neural progenitor cells (NPC), neural precursor cells (Precursor) and forebrain neurons (Neurons). (B) Immunoblot of Cav-1 isolated from Cav-1 WT iPSCs, NPC, Precursor and 10 weeks old Neurons. (C) Quantification of Cav-1 normalized to GAPDH expression and then normalized to Cav-1 WT iPSCs (n=3 replicates). (D) Immunoblots of Cav-1, β-III-Tubulin and GAPDH from iPSCs derived Cav-1 KO and Cav-1 WT NPCs, Precursor cells and 10 weeks old neurons. (E-F) Quantification of Cav-1 and β-III-Tubulin normalized to GAPDH expression in iPSCs derived Cav-1 KO and Cav-1 WT (n=3 replicates). (G) Representative confocal images of DCX (red) and DAPI (blue) positive cells in the iPSCs derived Cav-1 KO and Cav-1 WT NPCs, precursor cells and 5 weeks old neurons. Scale bar, 100 μm. (H) Representative confocal images of β-III-Tubulin (red) and DAPI (blue) positive cells in the iPSCs derived Cav-1 KO and Cav-1 WT NPCs, precursor cells and 5 weeks old neurons. Scale bar, 100 μm. (I) Quantification of DCX^+^DAPI^+^ cells iPSCs derived Cav-1 KO and Cav-1 WT NPCs, precursor cells and 5 weeks old neurons. n=9 images per group. (J) Quantification of β-III-Tubulin^+^DAPI^+^ cells iPSCs derived Cav-1 KO and Cav-1 WT NPCs, precursor cells and 5 weeks old neurons. n=9 images per group. (K) EdU cell proliferation assay in iPSCs derived Cav-1 WT and Cav-1 KO neural progenitor cells. Yellow arrowheads indicate EdU^+^DAPI^+^ cells. Scale bar, 50 µm. (L) Quantification of the percentage of EdU^+^ cells to the total DAPI comparing iPSCs derived Cav-1 WT and Cav-1 KO NPCs. Data represented as mean ± SEM. Data analyzed by unpaired two-tailed Student’s t-test (E, F, L), one-way ANOVA with Tukey multiple comparisons correction (C), two-way ANOVA with Sidak’s multiple comparisons correction (E, F, I, J). ns p > 0.05, *p < 0.05, **p < 0.01, ***p < 0.001, and ****p < 0.0001.

Cav-1 displayed a cell-type-specific, biphasic expression pattern during neuronal differentiation, with peak levels in NPCs and mature neurons (**Figure 5 B–C**). Cav-1 expression was low in iPSCs, increased in NPCs, and significantly decreased in precursor cells. Following differentiation, Cav-1 expression rose ∼6-fold in 10-week-old neurons compared to precursors. Cav-1 levels were also significantly higher in mature neurons than in iPSCs, NPCs, or precursors, suggesting a cell-autonomous role throughout differentiation. As expected, Cav-1 was expressed at all stages in control cells but was absent in Cav-1 KO cells, validating the model (**Figure 5 D–E**). Neuronal differentiation progression was further assessed by quantifying βIII-Tubulin expression (**Figure 5 D, F**).

Next, we asked whether Cav-1 similarly regulates neurogenesis in human NPCs, as observed in primary hippocampal NSPCs from Cav-1 WT and KO mice. DCX expression, assessed by immunocytochemistry in human NPCs, precursor cells, and 5-week-old neurons, was significantly higher in Cav-1 KO precursor cells than in WT controls (**Figure 5 G, I**). Cav-1 KO precursor cells and neurons showed significantly elevated β-III-tubulin expression, confirming enhanced neuronal differentiation relative to controls (**Figure 5 H, J**). Supporting our mouse data, Cav-1 KO iPSC-derived NPCs had significantly fewer EdU⁺ cells compared to WT, indicating reduced proliferation (**Figure 5 K-L**). Taken together, findings from iPSCs derived human NPCs and in vivo quantification of NSPC in mice confirmed that cell autonomous loss of Cav-1 halts cell proliferation while enhancing neuron differentiation in AHN.

### Rescue of Cav-1 expression in iPSCs-derived NPCs lacking Cav-1 restores mitochondrial morphology

To further assess the mechanism of Cav-1 as a direct regulator of mitochondria morphology, we used a lentivirus expressing Cav-1 and GFP (LV-Cav-1-p2A-GFP; LV Cav-1) or a GFP control (LV-GFP) to restore Cav-1 protein expression in our iPSCs model (**Figure 6 A**). Consistent with our proteomic and mouse NPCs model, we observed Cav-1 co-localization with the mitochondrial outer membrane marker TOM20 in iPSCs derived NPCs Cav-1 Ctrl and restoration of Cav-1 expression in Cav-1 KO NPCs using confocal microscopy (**Figure 6 B**). Next we performed live imaging using TMRM to examine rescue of mitochondria morphology (**Figure 6 C**). Consistent with our mouse NPC model, iPSCs-derived Cav-1 KO NPCs displayed significantly increased mitochondrial area, perimeter, and branch length, along with reduced circularity indicating a more elongated morphology compared to WT NPCs (**Figure 6 D–G**). Importantly, restoration of Cav-1 expression in the Cav-1 KO NPCs significantly reduced mitochondrial area, perimeter, and branch length, while increasing circularity compared to LV-GFP controls, confirming that Cav-1 regulates mitochondrial morphology in NPCs.

**Figure 6.**
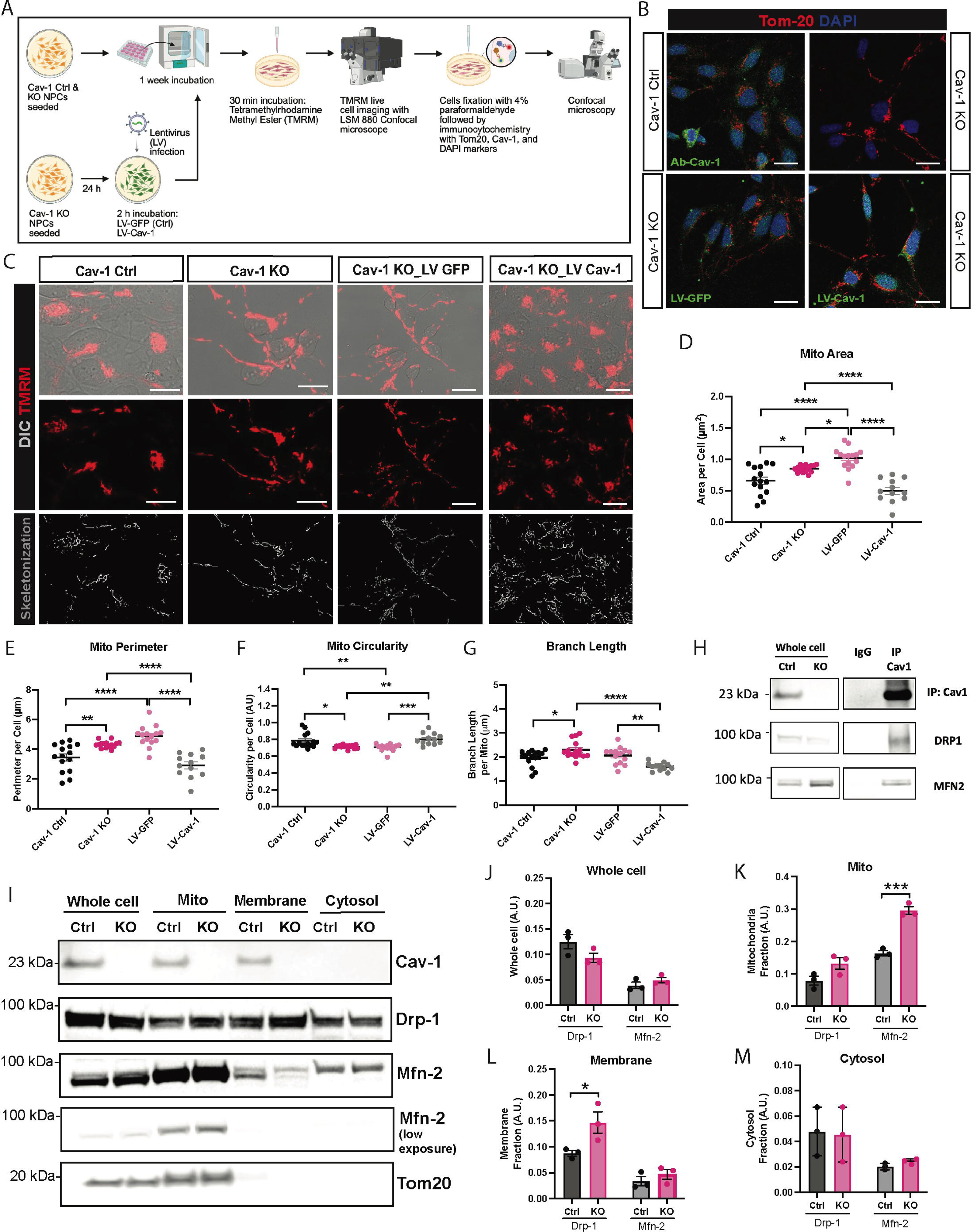
Cav-1 rescue restores mitochondrial morphology and dynamics via interaction with Mfn2 and Drp1 in iPSCs derived NPCs. (A) Live imaging experimental schematic representation with (Cav-1 rescue) and without lentivirus (LV) infection to NPCs. (B) Representative confocal image of Caveolin-1 (Cav-1, Green), Mitochondrial import receptor subunit TOM20 homolog (Tom-20, green) and DAPI (blue) immunostaining in Ctrl and KO NPCs. Below left image is of the Cav-1 KO NPCs infected with LV-GFP (Ctrl; green) and below right is with LV-Cav-1 (green). Scale bar 50 μm. (C) Representative live-cell images of TMRM staining from Cav-1 Ctrl, Cav-1 KO, Cav-1 KO NPCs with LV-GFP, and LV-Cav-1. NPCs were incubated with 50 nM TMRM for 30 min followed by confocal microscopy visualization. Mitochondria were skeletonized in ImageJ. Scale bar, 50 μm. (D-F) Quantification of mitochondria area per cell, mitochondria perimeter per cell, and circularity per cell in Cav-1 Ctrl, Cav-1 KO, Cav-1 KO LV-GFP, Cav-1 KO LV-Cav-1 NPCs. N=25 cells per group. (G) Quantification of skeletonized mitochondria branch length per mitochondria in Cav-1 Ctrl, Cav-1 KO, Cav-1 KO LV-GFP, Cav-1 KO LV-Cav-1 NPCs. N=25 cells per group. (H) Association of Cav-1 with MFN2 and DRP1. Cav-1 Ctrl NPCs lysates were immunoprecipitated with a Cav-1 polyclonal antibody and subjected to Western blot analysis with Cav-1, MFN2 and DRP1 Abs to assess co-IP. (I) Immunoblot of Cav-1, Drp-1, mitofusion-2 (Mfn-2) and Tom-20 in total cell lysate (Whole cell), mitochondrial (Mito) enriched, membrane, and cytosol fraction of Cav-1 Ctrl and Cav-1 KO NPCs. (J-M) Normalization of Cav-1 KO Drp-1 and Mfn-2 protein expression in whole cell, mito, membrane, and cytosol enriched fractions to Cav-1 Ctrl expression. Data representative of n=3, 6-well plate per genotype. Data represented as mean ± SEM. Data analysed by unpaired two-tailed Student’s t-test except in (D-G) was analyzed by two-way ANOVA with Tukey’s multiple comparisons correction. *p < 0.05, **p < 0.01, ***p < 0.001 and ****p < 0.0001.

To further investigate the mechanism by which Cav-1 regulates mitochondrial morphology, we assessed its interaction with key mitochondrial dynamics regulators: mitofusin 2 (Mfn2) and dynamin-related protein 1 (Drp1). Mfn2 promotes mitochondrial fusion, enabling the elongation and interconnection of mitochondrial networks, while Drp1 mediates mitochondrial fission, leading to mitochondrial fragmentation and distribution. Co-immunoprecipitation confirmed that both Mfn2 and Drp1 directly interact with Cav-1 (**Figure 6 H**). Next, alterations in the mitochondrial localization of Drp1 and Mfn2 in the absence of Cav-1 were evaluated through subcellular fractionation of Cav-1 WT and Cav-1 KO NPCs (**Figure 6 I-M**). Western blot analysis revealed a significant increase in Mfn2 levels in the mitochondrial fraction with no change in Drp-1 levels as result of Cav-1 loss (**Figure 6 K).** Interestingly, a significant increase in membrane Drp-1 levels were found in the Cav-1 KO NPCs compared to control (**Figure 6 L).** No change in total expression levels or cytosol expression levels of Mfn2 or Drp1 were found in Cav-1 KO NPCs compared to controls (**Figure 6 J, M**). Together, our findings suggest that Cav-1 regulates mitochondrial morphology by modulating the subcellular localization of fusion-fission-linked proteins and therefore, Cav-1 loss promotes neuronal lineage commitment by enhancing mitochondrial fusion in NPCs.

### Restoring Cav-1 levels reverses neuronal differentiation phenotype

To examine whether reinstating Cav-1 in Cav-1KO NPCs would reverse enhanced neurogenesis, Cav-1 in Cav-1 KO NPCs were infected with LV-GFP (control) or LV-Cav-1. Examination of Cav-1 expression in protein lysate of these cells across neural differentiation stages validated successful transduction procedure (**Figure 7A, B)**. Notably DCX protein levels is significantly reduced in precursors with Cav-1 overexpression, indicating normalization of enhanced neuronal lineage commitment at this stage (**Figure 7C**). Strong trends toward decreased β-III-Tubulin, were observed in Cav-1-overexpressing lines, though these differences do not reach statistical significance at precursor and neurons stage (**Figure 7D**). Together with **Figure 6** data, these results suggest that Cav-1 regulates neurogenesis by modulating mitochondrial remodeling (**Figure 7E).**

**Figure 7.**
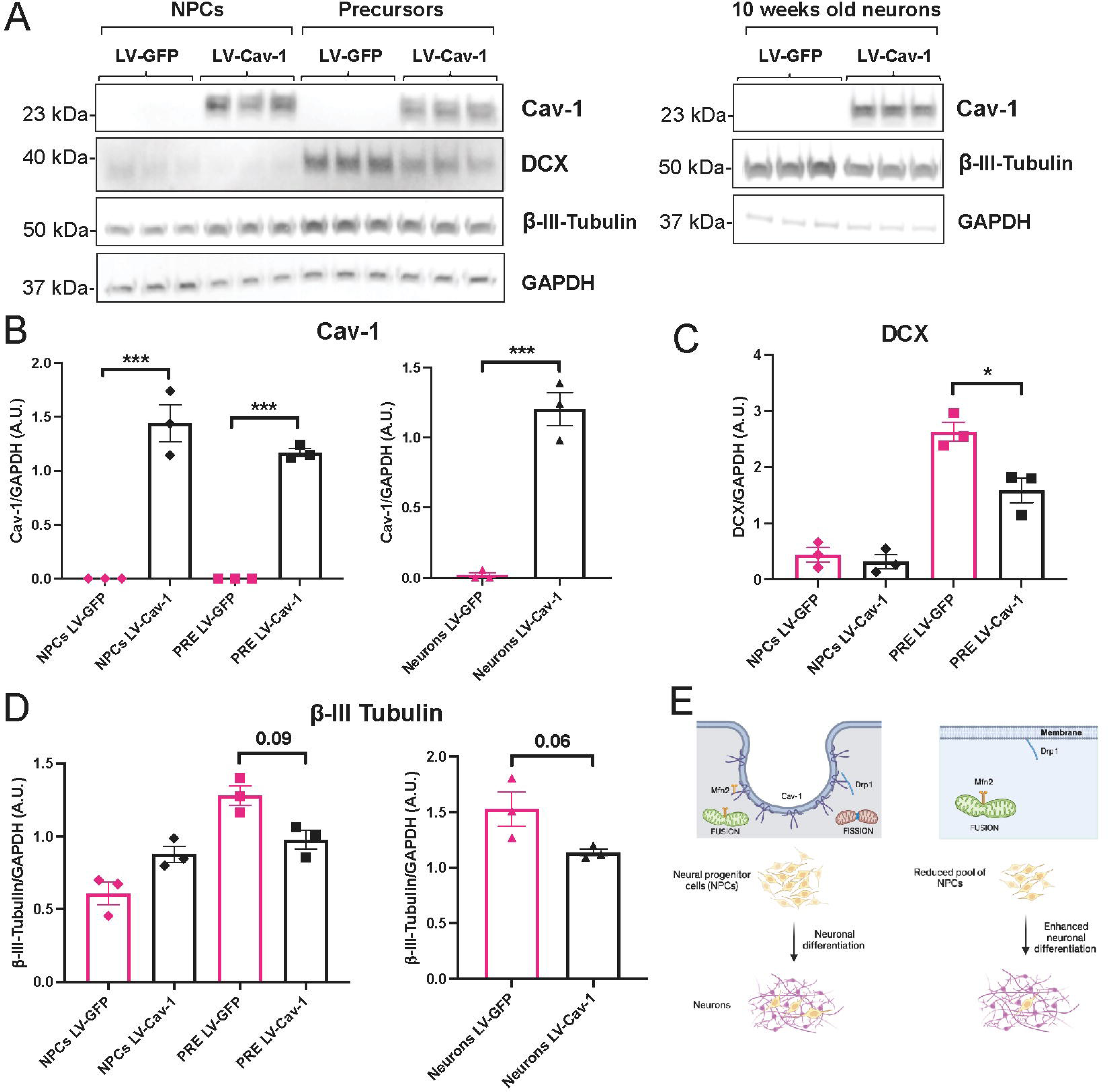
Restoring Cav-1 levels normalizes neuronal differentiation process. (A) Representative Western blots showing expression levels of Caveolin-1 (Cav-1), DCX, β-III-Tubulin, and GAPDH across neural progenitor cells (NPCs), precursors (PRE), and 10-week-old neurons, transduced with either LentiVirus (LV)-GFP (control) or LV-Cav-1. (B-D) Quantifications of Cav-1, DCX, and β-III-Tubulin normalized to GAPDH expression in iPSCs derived Cav-1 KO NPCs, transduced with either LV-GFP or LV-Cav-1, NPCs-induced differentiation into precursor and neurons. Data representative of n=3, 6-well plate per treatment. Data represented as mean ± SEM. Data analyzed by unpaired two-tailed Student’s t-test (neurons), and one-way ANOVA with Tukey multiple comparisons correction (NPCs and PRE). *p < 0.05, ***p < 0.001. (E) Schematic illustrating Cav-1’s role in facilitating mitochondrial fusion-fission dynamics in NPCs, thereby regulating neuronal differentiation.

## Discussion

This study revealed several key insights. First, Cav-1 deletion in mouse NSPCs reduced NSPC proliferation and increased the number of immature neurons in the DG, suggesting that Cav-1 acts as a negative regulator of neuronal differentiation. This shift in cell fate was associated with enhanced context discrimination memory and greater dendritic volume in newborn neurons, indicating functional relevance. Next, to extend these findings to human neurogenesis, we used iPSCs-derived NPCs and found that, similarly to the mouse, Cav-1 deletion impaired proliferation and promoted neuronal differentiation, supporting a conserved role for Cav-1 in regulating neurogenesis. Finally, we identified Cav-1 as a novel regulator of mitochondrial dynamics in neurogenesis. Cav-1 loss in human NPCs led to elongated mitochondria, consistent with increased fusion, which was reversed by Cav-1 restoration. Mechanistically, Cav-1 directly bound Mfn2 and Drp1, key mediators of fusion and fission, respectively. In Cav-1 KO NPCs, mfn2 was enriched in mitochondrial fractions, while Drp1 remained in membrane fraction, indicating that Cav-1 supports balanced mitochondrial dynamics by facilitating proper protein localization. Together, these findings establish Cav-1 as a critical coordinator of neurogenesis and mitochondrial function.

The balance between quiescence, proliferation, and differentiation of neural stem cells (NSCs) is essential for maintaining hippocampal neurogenesis. While NSCs integrate both extrinsic and intrinsic signals to determine their fate (Urban et al., 2019), our findings identify Cav-1 as a key intrinsic regulator that suppresses neuronal differentiation. Although Cav-1 is enriched in endothelial cells and may exert non-cell autonomous effects, our data show that its deletion in NSCs leads to a reduction in NSPC proliferation and a marked increase in immature neurons, suggesting a shift toward neuronal fate commitment. This was accompanied by enhanced dendritic volume in newborn neurons, a hallmark of functional maturation and integration into hippocampal circuits.

Dendritic growth is critical for synaptic integration, input specificity, and neuronal survival. In the dentate gyrus, adult-born granule cells must extend complex dendritic trees to reach the molecular layer and receive afferent input from the entorhinal cortex. Increased dendritic volume and complexity have been correlated with enhanced synaptic input, improved synaptic plasticity, and stronger contributions to hippocampal-dependent learning tasks (Zhao et al., 2006; Toni et al., 2007). Thus, the observed dendritic phenotype in iNSC Cav-1 KO mice likely enhances the functional relevance of increased neuronal differentiation by potentially facilitating synaptic incorporation and circuit maturation.

Recent studies suggest that mitochondrial morphology is tightly linked to stem cell identity and cell fate, with mitochondrial fusion promoting neuronal differentiation (Khacho et al., 2016; Khacho and Slack, 2018; Coelho et al., 2022). NSC in the adult DG exhibit fragmented and globular mitochondria that become more elongated as cells progress into committed NPCs and immature neurons (Beckervordersandforth et al., 2017). *In vivo* and *in vitro* studies show that when NSPC differentiate into neurons a metabolic switch occurs from glycolysis to OXPHOS (Zheng et al., 2016; Beckervordersandforth et al., 2017). Moreover, mitochondrial fission-fusion dynamics, orchestrated by Drp1 and Mfn2, are increasingly recognized as essential for newborn neuron plasticity and maturation (Kochan et al., 2024).

Our findings add to the growing body of evidence implicating mitochondrial dynamics in neural stem cell fate by identifying Cav-1 as a novel regulator of mitochondrial morphology in neurogenesis. We show that Cav-1 physically interacts with both Drp1 and Mfn2, suggesting it may serve as a scaffold to coordinate their subcellular localization. Deletion of Cav-1 in both mouse NSPCs and human NPCs led to elongated mitochondrial morphology, consistent with enhanced mitochondrial fusion, and increased mitochondrial velocity and displacement in mouse NSPCs. In human Cav-1 KO NPCs, we observed a significant redistribution of Mfn2 to the mitochondria and Drp1 to the plasma membrane, without changes in total protein levels, indicating a spatial rather than transcriptional regulation. Notably, prior studies have shown that loss of Cav-1 or mutation at its phosphorylation site disrupts its interaction with Mfn2, thereby permitting Mfn2 translocation to the mitochondrial outer membrane and thus, alter cell migration patterns (Jiang et al., 2022). Importantly, re-expression of Cav-1 in Cav-1 KO NPCs restored mitochondrial morphology, further supporting its regulatory role in mitochondrial dynamics. This shift toward a fused mitochondrial state aligns with our observed increase in neuronal differentiation in Cav-1-deficient models, suggesting that Cav-1 normally acts to restrain lineage commitment by maintaining mitochondrial dynamics.

Our previous studies further underscore the relevance of Cav-1 function in disease states associated with impaired neurogenesis (Bonds et al., 2019; Bonds et al., 2020). We found that Cav-1 levels were significantly reduced in the hippocampi of patients with type 2 diabetes mellitus (T2DM) and inversely correlated with β-amyloid (Aβ) levels. Similarly, in the db/db diabetic mouse model, Cav-1 expression was diminished alongside elevated levels of amyloid precursor protein (APP) and hyperphosphorylated tau—hallmarks of Alzheimer’s disease (AD) pathology (Bonds et al., 2019). Notably, these mice also exhibited impaired hippocampal neurogenesis (Bonds et al., 2020), consistent with our current findings that link Cav-1 deficiency to altered proliferation of NSC in the DG. Furthermore, we previously demonstrated that reduced endothelial Cav-1 expression contributes to insulin signaling deficits observed in T2DM (Shetti et al., 2023). Given the link between mitochondrial dysfunction and both T2DM and AD (Carvalho and Moreira, 2023), these findings suggest that Cav-1 may be a contributing factor in the neurogenesis and cognitive impairments associated with these disorders.

In conclusion, this study identifies Cav-1 as a novel intrinsic regulator of adult hippocampal neurogenesis that exerts its effects, in part, by governing mitochondrial dynamics. Cav-1 directly interacts with Mfn-2 and Drp-1, supporting the recruitment of these proteins to mitochondria and maintaining balanced fusion and fission. Loss of Cav-1 favors mitochondrial fusion, accelerates NSPC differentiation, and enhances neurogenesis-dependent cognitive function. These findings broaden our understanding of the molecular interplay between membrane scaffolding proteins and mitochondrial dynamics in the regulation of adult neurogenesis.

## Materials and Methods

### Mouse Models

All mouse experiments were approved by the University of Illinois at Chicago Institutional Animal Care and Use Committee. Mice were housed on a 12-hour light/dark cycle and provided food and water ad libitum. C57Bl/6J wildtype (WT) and global Cav-1 knockout (gCav-1 KO) used in experiments were from Jackson Laboratories (Strain #:000664 and Strain #: 007083, respectively). Inducible NSPC specific-Cav-1 knockout mice (NestinCre^ERT2/+^;Cav-1^fl/fl^) were generated by crossing NestinCre^ERT2/+^ mice (Sahay et al., 2011; Mishra et al., 2022) with Cav-1^fl/fl^ mice (Cao et al., 2003; Oliveira et al., 2017). The NestinCre^ERT2/+^ transgene was maintained as hemizygous and the Cav-1^fl/fl^ homozygous. Male mice were used in experiments unless otherwise stated in text.

### Tamoxifen Injections

Tamoxifen (TAM, Sigma-Aldrich) was dissolved in corn oil (Sigma-Aldrich) at 20 mg/mL at 37°C and then stored at 4°C for 5 days. To induce recombination, 4–5-week-old mice were intraperitoneally injected at a dose of 130 mg/kg TAM or equal volume of corn oil once a day for 5 consecutive days.

### Viral Vectors

The Retrovirus/Moloney Murine Leukemia Virus (MMLV) construct carrying GFP (RV-GFP) was generously provided by Dr. Jenny Hsieh (University of Texas at San Antonio, TX). The viral particles were packaged as previously described (Lybrand et al., 2021). The packaging process was carried out at the Vector Core Facility of the University of Illinois Chicago (UIC) under the guidance of the Hsieh lab. The viral titer was 1.67 × 10L TU/ml for this study.

### Mouse NSPC Isolation and Culture

Primary hippocampal NSPCs were isolated from 6–8-week-old mice similarly as described (Bonds et al., 2020; Ahmed et al., 2021). Hippocampi were dissected in ice-cold HBSS and pooled from 4-6 mice. Hippocampi were transferred to a tissue culture hood, minced using a sterile scalpel until no visible pieces remained (3-5 mins) and transferred to 3-5 mL of warm culture media (DMEM/F-12 with 20 mM KCl, 2 μg/mL heparin, 1% penicillin–streptomycin, 20% B27 supplement, 10% N2 supplement). Tissue was spun at 200g for 2 mins and dissociated with 0.1% Trypsin-EDTA (diluted in DMEM/F12) at 37°C for 10 mins. After incubation, tissue was triturated 5 times with a P1000 pipette. Next, 3 mL of trypsin-inhibitor (0.139 mg/mL and 1U/mL DNase I in HBSS-/-) was added, triturated with a P1000 pipette 5 times and centrifuged at 300g for 5 mins. The cell pellet was singly resuspended in 1 mL of culture media by pipetting an additional 25 times with a P1000 pipette followed by filtration through a 40 μm cell strainer and filter washed with 15-20 mL of culture media. Cells were centrifuged 300g for 5 mins and resuspended in 1 mL of culture media plus growth factors (20 ng/mL EGF and 10 ng/mL bFGF) by pipetting 10 times with a P1000 pipette. Cells were plated in a 24-well plate (1 well per mouse) containing 2 mL of culture media plus growth factors. Half media changes were preformed every 48 hrs for 7-10 days with growth factors added to culture media immediately prior to use.

After the first 7-10 days, cells were collected and centrifuged 300g for 5 mins. The cell pellet was singly dissociated by pipetting 20 times with a P1000 pipette in 1 mL of culture media containing growth factors. The cell suspension was filtered through a 40 μm cell strainer and filter washed with 15-20 mL of culture media. Cells were centrifuged 300g for 5 mins, resuspended in 1 mL of culture media plus growth factors and plated in 2-3 wells of 6-well plate containing 3 mL of culture media plus growth factors per well. Half media changes were preformed every 48 hrs until ∼100 μm in diameter neurospheres are formed (7-10 days). All experiments were performed using NSPCs between passage numbers 3 to 8, where neurospheres were dissociated using Accutase (StemCell Technologies) at 37°C for 7 mins followed by the addition of 5 mL culture media and centrifuged 300g for 5 mins. Cells were counted and plated at 10,000 cells/cm^2^ for floating neurosphere cultures. For experiments requiring monolayers, singly dissociated NSPCs were grown on poly-L-orthinine/laminin (PLO/laminin) coated plastic dishes or acid washed coverslips. Briefly, plastic dishes and coverslips were incubated at 37°C with 15 μg/mL PLO diluted in 1X PBS for 24-48 hr, washed with 1X PBS and incubated at 37°C with 10 μg/mL laminin in 1X PBS for 24 hr. PLO/laminin coated dishes or coverslips were washed 1 time with 1X PBS prior to plating of cells.

### Human iPSCs Culture, NPC differentiation and Neuron Maturation

CRISPR-sham- and CRISPR/Cas9-edited induced pluripotent stem cell lines (Cav-1WT and Cav-1KO iPSCs, n = 4) were generated by Synthego. These iPSCs lines were derived from adults with no known trauma or genetic dispositions. One vial from the master stocks was thawed at 37°C and were plated on matrigel-coated plates (MG, Corning) with mTeSR plus media and Y-27632 (ROCK inhibitor) used at a final concentration of 10μM. The ROCK inhibitor was added during the initial 24 h medium which was replaced daily. Individual colonies were mechanically dissected and were passaged weekly. All reagents were purchased from STEMCELL Technologies unless otherwise specified.

The neural differentiation protocol was followed from STEMCELL Technologies and the NPCs, precursor cells, 10 weeks old neurons were validated by ICC staining. Refer figure 5A. for the differentiation timeline. STEMdiff Neural Induction Medium + SMADi was used to generate CNS-type NPCs from hiPSCs using embryoid body (EB) protocol. STEMdiff Neural Progenitor Medium was used to culture and passage neural progenitor cells. STEMdiff™ Forebrain Neuron Differentiation Kit was used to generate neuronal precursors from NPCs. The neuronal precursors generated were matured using STEMdiff™ Forebrain Neuron Maturation Kit to produce forebrain-type neurons.

Confluent iPSCs were dissociated using Gentle Cell Dissociation Reagent and plated at 3×L10^6^ cells/well in neural induction medium (NIM) with SMADi of the AggreWell^TM^800 plate, treated with Anti-Adherence Rinsing Solution. This resulted in 10,000 cells/microwell. Y-27632 was added to STEMdiff^TM^ Neural Induction Medium + SMADi to obtain a final concentration of 10μM for the first 24h only. NIM was partially replaced daily for 5 days, after which EBs were harvested from a single well of an AggreWell^TM^800 plate and plated onto a tissue culture-treated 6-well plate coated with PLO/laminin for the formation of neural rosettes. After a week, neural rosettes were selected using STEMdiff^TM^ Neural Rosette Selection Reagent and replated on PLO/Laminin coated 6-well plate for the generation for NPCs differentiation. Cells were either expanded for banking or kept for neuronal differentiation. One fifth of cells were retained at each passage for marker analysis. Cells were expanded up to 4 passages.

NPCs were cultured for a week in neural progenitor medium. After seven days, cells were dissociated using accutase, and re-plated in Forebrain Neuron Differentiation media for neural precursor cells differentiation for a week. The neuronal precursors generated were matured using STEMdiff™ Forebrain Neuron Maturation Kit to produce a mixed population of excitatory and inhibitory forebrain-type neurons (≥ 90% class III β-tubulin-positive neurons). Cell culture at all stages was carried out on tissue culture plastic well plates and for imaging, 24 well glass bottom plates were used.

### Cav-1 Lentivirus Transduction

A custom-designed lentiviral vector was purchased from VectorBuilder. The construct was designed to express Cav-1 and GFP separated by a self-cleaving P2A linker to allow equimolar co-expression from a single transcript under the control of the Nestin promoter (Vector ID: VB200519-1185asg). The control lentiviral construct was designed to express GFP under the control of the Nestin promotor (VB200803-1257vht). The viral titer was >2×10^9^ TU/ml for this study.

NPCs were cultured as described above. Approximately 50,000 NPCs were seeded per well in 24-well glass bottom plates. Approximately 80% confluent NPCs were transduced using produced virus particles at multiplicity of infection (MOI) of 10. The medium was changed after 2 h and the cells were analyzed 5 days post-transduction, after TMRM incubation for 30 min, using a confocal microscope.

Similarly, for the neuronal differentiation rescue experiment, approximately 1.2 x 10^6^ NPCs were seeded per well in a 6-well plate. After 24 hours, cells were transduced with lentivirus (LV) carrying either GFP or Cav-1 for 2 hours, followed by a medium change. Neuronal differentiation was then carried out as described previously. NPCs were maintained in neural progenitor medium for one week, differentiated into precursor cells using Forebrain Neuron Differentiation Medium for another week, and subsequently matured using the STEMdiff™ Forebrain Neuron Maturation Kit for up to 10 weeks.

### BrdU and EdU Labeling

For the 12-day *in vivo* experiments, 5-bromo-2’-deoxyuridine (BrdU, Sigma) was prepared fresh daily by dissolving BrdU at 20 mg/mL in 1X PBS at 37°C and then sterile filtered via 0.22µm syringe filter. Mice were intraperitoneally injected at a dose of 100 mg/kg once daily for 12 consecutive days and tissue collected 24 hr after the last injection. Mice were sacrificed and tissue collected 4 weeks after the last injection. For *in vitro* BrdU and EdU experiments, NSPCs were seeded on PLO/laminin coated coverslips at a density of 50,000 cells/cm^2^ and incubated overnight in culture media containing growth factors. A 5 mM EdU stock was prepared in DMSO and stored at −20°C until use. EdU stock solutions were diluted to 5 μM in culture media, sterile filtered via 0. 22µm syringe filter and growth factors added immediately prior to the start of the experiment.

### Clonogenic Proliferation Assay

Hippocampal NSPCs were singly dissociated and plated as floating cultures in 96-well plates at 1000 cells per well. Cells were grown in culture media containing growth factors (20 ng/mL EGF and 10 ng/mL bFGF) and growth factors were added to media every 48 hr. On day 6, cultures were imaged using a Keyence BZ-X800 followed by dissociation with Accutase to count the number of cells per well. The number and diameter of neurospheres (clones) per well was quantified using ImageJ.

### Differentiation Assay

Hippocampal NSPCs were seeded on PLO/laminin coated plastic dishes or coverslips at a density of 50,000 cells/cm^2^. Cells were grown in culture media containing growth factors (20 ng/mL EGF and 10 ng/mL bFGF) overnight (8-12 hours). Media was then changed to differentiation media consisting of culture media (DMEM/F-12 with 20 mM KCl, 2 μg/mL heparin, 1% penicillin–streptomycin, 20% B27 supplement, 10% N2 supplement) with 1 μM retinoic acid (Sigma) and 5 μM forskolin (Sigma). Half of the media was changed every 48-72 hours. Cells were fixed or collected after 3 or 7 days of differentiation.

### Immunohistochemistry and Immunocytochemistry

Mice were transcardially perfused with ice cold PBS followed by 4% PFA in 1X PBS. Brains were post-fixed for 24 hr in 4% PFA followed by immersion in 10% sucrose in 1X PBS for 24 hr, 20% sucrose in 1X PBS for 24 hr and 30% sucrose 1X PBS for 24 hr. Brains were sectioned at 40 μm using a sliding microtome and floating coronal sections were stored in cryoprotectant consisting of glycerol (20% v/v) and ethylene glycol (24% v/v) in 1X PBS at −20°C. Sections were washed with 1X PBS, incubated with 1% sodium borohydride in 1X PBS for 10 mins at RT and washed 3 times with 1X PBS for 10 min per wash. For antibodies requiring heat induced antigen retrieval, sections were incubated in 10 mM sodium citrate buffer containing 0.05% Tween 20, pH 6.0 at 99°C for 15 mins in a vegetable steamer. For BrdU immunodetection, sections were washed with 1X PBS, treated in pre-warmed 1N HCl for 40 min at 37°C followed by incubation with 0.1 M sodium borate buffer pH 8.5 for 10 min at RT. Following, antigen retrieval or HCl pre-treatment, sections were washed 3 times with 1X PBS for 10 min per wash. Sections were blocked in 1X PBS containing 0.3 M Glycine, 0.2% Triton X-100 and 5% normal donkey serum (NDS) for 1 hr at RT. After blocking, sections were incubated with primary antibodies diluted in block solution for 48 hr at 4°C. Primary antibodies used were, Rat anti-BrdU (ab6326), Rabbit anti-Cav-1 (3267), Rabbit anti-DCX (4604), Mouse anti-DCX (sc-271390), Guinea pig anti-DCX (AB2263), Rabbit anti-GFAP (Z0334), Rat anti-GFAP (13-0300), Rabbit anti-MCM2 (4007), Chicken anti-Nestin (AvesLab), Rabbit anti-NeuN (ab104225), Mouse anti-NeuN (ab104224), Rabbit anti-Sox2 (ab97959), Goat anti-Sox2 (AF2018), Rat anti-Tbr2 (14-4875-82). After primary incubation, sections were washed 3 times in PBS containing 0.1% Tween-20 (PBST) for 15 mins per wash. Sections were incubated with secondary antibodies diluted in block solution for 1.5-2 hr at RT. Secondary donkey antibodies used were ordered from Jackson ImmunoResearch Labs. Sections were washed 3 times in PBST for 15 mins per wash, counterstained with DAPI, and mounted on Superfrost Plus slides (ThermoFisher) with Prolong Gold Antifade Mountant (Invitrogen).

For immunocytochemistry (ICC), culture media was removed, coverslips containing cells were washed two times with 1X PBS and fixed in 4% PFA in 1X PBS for 20 min at RT. Cells were washed two times with 1X PBS and blocked in 1X PBS containing 0.3 M Glycine, 0.2% Triton X-100 and 5% NDS for 30 minutes at RT. For EdU detection, cells were processed according to the Click-iT EdU Cell Proliferation Kit (Invitrogen) instructions prior to blocking step.

For BrdU immunodetection, cells were treated in pre-warmed 1N HCl for 40 min at 37°C followed by incubation with 0.1 M sodium borate buffer pH 8.5 for 10 min at RT prior to blocking step. Primary antibodies were diluted in blocking solution and incubated 24 hr at 4°C. Primary antibodies used were, Rabbit anti-DCX (ab18723) and Mouse anti-βIII-tubulin (MA1-118). After primary incubation, cells were washed 3 times in PBST for 5 mins per wash and incubated with secondary antibodies diluted in blocking solution for 45 min at RT. Cells were then washed 3 times in PBST for 5 mins per wash, counterstained with DAPI and mounted with Prolong Gold Antifade Mountant (Invitrogen). Images were acquired at 40x or 63x magnification using confocal microscopy (Zeiss LSM 710).

### GFP Retrovirus Hippocampal Stereotaxic Injection

The Retrovirus/Moloney Murine Leukemia Virus (MMLV) construct carrying GFP (RV-GFP) was generously provided by Dr. Jenny Hsieh (University of Texas at San Antonio, TX). The viral particles were packaged as previously described (Lybrand et al., 2021). The packaging process was carried out at the Vector Core Facility of the University of Illinois Chicago (UIC) under the guidance of the Hsieh lab. The viral titer was 1.67 × 10L TU/ml for this study.

Stereotaxic surgeries were conducted on mice aged 4 to 4.5 months (16-18 weeks). Anesthesia was induced using a 2.5% isoflurane and 100% oxygen mixture, and buprenorphine (0.01 mg/kg) was administered preoperatively for analgesia. A total of 1 μl of retrovirus was bilaterally injected into the dentate gyrus (DG) at three stereotaxic coordinates: AP = −1.50 mm, ML = ±0.60 mm, and DV = −2.25 mm; AP = −2.00 mm, ML = ±1.30 mm, and DV = −2.15 mm; and AP = −2.55 mm, ML = ±1.90 mm, and DV = −2.20 mm. Injections were delivered at a rate of 70 nl/min using a 1 μl Hamilton microsyringe. The needle was positioned at the target site for 5 minutes before injection and remained in place for an additional 10 minute post-injection to prevent viral backflow. Mice were sacrificed 28 days after the procedure, and injection sites were confirmed through histological and immunohistochemical analysis.

### Neuronal Reconstruction, Sholl and Dendritic Analysis

Imaging of new neurons (labeled with RV-GFP) in the dentate gyrus of CORN and TAM injected NestinCre^ERT2/+^; CAV-1^fl/fl^ was accomplished with Neurolucida 360 (MBF Bioscience, Inc.) and a Zeiss LSM 710 confocal microscope. Neurons were imaged under 20X magnification and centralized to include the soma and all neurites. Once image was acquired, Neurolucida would be used to create a 3D tracing of the cell. Briefly, soma tracing would be used to create and automated tracing of the cell body. Once the soma was finished, dendritic tree tracing was used, using the user guide mode (a semi-automated setting where Neurolucida automatically detects the thickness of the processes while their path is being guided by the user) to create the final tracing of the cell.

For the Sholl analysis, the Neurolucida explorer software (MBF Bioscience, Inc.) was used to report the number of intersections per pre-determined radius for each neuronal trace. Starting at a radius of 10 μm from the center of the soma, a continuous shell of concentric circles with 10μm between them was used to determine the number of intersections per radius. For determining the total surface area (μm^2^) and volume (μm^3^) of the dendritic tree, Neurolucida explorer software was used. Using the neuron summary function in the branched structure analysis of the software, a total surface area and volume for all the dendritic branches of the cell were analyzed. For the Sholl analysis, the number of intersections per radius was analyzed using a mixed effects model with the Geisser-Greenhouse correction followed by Bonferroni post-hoc test for multiple comparisons.

### Transmission Electron Microscopy

Cells were washed 3 times with 1X PBS and centrifuged at 300g for 5 min to form pellets. Cell pellets were fixed in 1.6% glutaraldehyde in 100 mM sodium phosphate, pH 7.4 for 1 hr at room temperature as similarly described(Sandhu JK, 2021). Samples were post-fixed with 1% osmium tetroxide for 1 hr and dehydrated using an ascending series of ethanol (through 100% absolute). Sample were then embedded in LX112 epoxy resin and polymerized at 60°C for 3 days. Ultrathin sections (∼75 nm) were collected onto copper grids and stained with uranyl acetate and lead citrate, respectively. Specimens were examined using a JEOL JEM-1400F transmission electron microscope at 80 kV. Micrographs were acquired using an AMT Side-Mount Nano Sprint Model 1200S-B and Biosprint 12M-B cameras, loaded with AMT Imaging software V.7.0.1.

### RNA Isolation

To isolate RNA, cells were washed 2 times with 1X PBS and RNA was isolated using a RNAeasy Plus Mini Kit (Qiagen).

### Quantitative Real-Time PCR

Quantitative real-time PCR (qRT-PCR) was used to measure RNA with Luna Universal One-Step RT-qPCR Kit (New England Biolabs) via CFX Connect Real-Time PCR Detection System (Bio-Rad). Target gene expression was normalized to gene expression of GAPDH or β-Actin.

### Western blotting

Cells were washed 3 times with 1X PBS and lysed on ice for 10 mins in RIPA buffer containing protease and phosphatase inhibitor cocktails (ThermoFisher). Lysed samples were sonicated on ice at 20% power 3 times for 15 s with 5 s rest between sonication. Samples were centrifuges at 10,000g for 15 mins to remove insoluble material and cellular debris. The supernatant was collected and protein concentration determined by BCA Protein Assay Kit (ThermoFisher). Samples were prepared in sample buffer consisting of NuPAGE LDS Sample Buffer and NuPAGE reducing agent followed by boiling for 5 mins at 95°C. Protein were separated on Bolt Bis-Tris Plus SDS-PAGE gels (Invitrogen) with MES SDS running buffer (Invitorgen) and transferred to 0.2 μm nitrocellulose membranes via the iBlot 2 Dry Blotting System (Invitrogen) Membranes were blocked for 1 hr at RT in 5% non-fat dry milk (milk) diluted in TBS containing 0.1% Tween-20 (TBST). Primary antibodies were diluted in 5% milk in TBST and membranes incubated overnight at 4°C. Primary antibodies used were, Rabbit-anti-caveolin-1 (3267), Mouse anti-βIII-tubulin (MA1-118), Rabbit-anti-TOM20 (42406S), Rabbit-anti-DRP1 (8570), and Mouse-anti-MFN2 (67487-1-Ig). Membranes were washed 3 times for 15 mins per wash with TBST followed by incubation with HRP conjugated secondary antibodies [Goat anti-mouse IgG-HRP (W402B), and Goat anti-rabbit IgG-HRP (W4011)] diluted in 5% milk in TBST for 2 hr at RT. Membranes were washed 3 times for 15 mins per wash and developed with ECL Super Signal Kit via Kodak RP X-OMAT developer or Azure Biosystems 300q Image Western Blot Imaging System. Band intensities were quantified in Fiji (NIH) from scanned images and total protein expression was normalized to GAPDH or β-Actin protein expression.

### Cell Quantification

For *in vivo* experiments, every sixth section of brain tissue was quantified using unbiased stereology (StereoInvestigator, MBF Biosciences). Under the optical fractionator workflow, contours of the DG were traced under 10x magnification and cells counted under 63x magnification. A 120 μm x 120 μm counting frame with a 2 μm guard zone on both sides of the section and a counting grid size determined by sampling 35% or 50% of the contour was used. The volume of DG (μm^3^) was determined by multiplying the area of contour by the measured mounted thickness of the section. Total cell counts were normalized to volume of DG (μm^3^) counted per mouse. Alternatively, for DCX and NeuN cell quantification, 30 μm z-stacks of the DG were acquired at 25x magnification using confocal microscopy (Zeiss LSM 710) and positive cells counted from maximum projection images in Fiji (NIH). For in vitro experiments, 10 μm z-stacks were acquired at 40x or 63x magnification using confocal microscopy (Zeiss LSM 710) and positive cells counted from maximum projection images in Fiji (NIH).

### Behavior Tests

All mice were handled 3-5 days for 2 min per mouse per day prior to the start of behavior testing.

### Elevated Plus Maze

The elevated plus maze (EPM) test was used to examine anxiety-like behavior. The EPM apparatus consisted of two open arms without walls and two closed arms with opaque walls. Mice were placed at the center of the apparatus facing an open arm and allowed to freely explore for 5 mins. The EPM apparatus was cleaned with 70% ethanol between mice. Video recordings were analyzed by Ethovision XT v16 software (Noldus) and the time spent in the open and closed arms as well as frequency of entries into the arms was calculated.

### Novel Object Location

The novel object location (NOL) test was used to examine spatial learning and memory behavior as similarly described in X. A 38 cm x 51 cm x 30 cm opaque white plastic chamber box with one short end containing a black circle wall print and the other short end containing a black vertical line wall print was utilized. On day 1 of the test, mice were habituated in the empty box for 10 mins followed by placement of 2 identical objects equal distance a part on the short end of the box containing the black circle wall print for 10 mins. Objects were removed for 5 min and then replaced in same locations for another 10 min session. Mice were placed back in home cages for 24 hrs. On day 2, mice were habituated in the empty box for 10 mins and tested by placing one of the objects in a novel location diagonally to the other one along the short end of the box with black vertical line wall print. Mice were allowed to explore the objects for 5 mins. Video was captured for every trial and exploration time with each object was manually scored blinded. Chambers were cleaned with 70% ethanol between mice. The discrimination index (DI) was calculated as DI = (T_N_-T_o_)/(T_n_+T_o_), where T_N_ is the exploration time with the object in the new location and T_o_ the exploration time with the object in the old location.

### Contextual Fear Discrimination

This contextual fear discrimination test was conducted similarly as described(Clemenson et al., 2014). All contexts consisted of 17.8 cm x 17.8 cm x 30.5 cm chamber housed in a sound isolation cubicle (Coulbourn Instruments). Context A consisted of two translucent plexiglass walls, two walls comprised of alternating black and white metal tiles and a stainless-steel rod floor. Context A also had a 28V exhaust fan. Context B had no fan and consisted of two walls comprised of alternating black and white metal tiles, two walls of black vertical line print and a stainless-steel rod floor. To test fear generalization, Context B was exchanged with Context C that had no fan, a circular wall insert made of up of a black and white plastic sheet and a black plastic floor covering. On day 1 of the test, mice were exposed to Context A for 10 mins. On day 2, mice were placed in Context A, received a 2s foot shock (0.7mA) immediately upon entry and remained in Context A for an additional 28s (a total of 30s in Context A). Thirty mins later, mice were placed back in Context A for a 3 min exposure trial followed by placement in Context B for 3 min exposure trial. On day 3 (24 hr post-shock), mice were placed in Context A for 3 mins or Context B for 3 mins in a counter-balanced order. For a less difficult discrimination test, mice followed the same sequence as stated but were placed in Context C instead of Context B on day 2 and 3. FreezeFrame software (Actimetrics) was used for video recording and analysis of freezing behavior in each context. Chamber walls and floor were cleaned with 70% ethanol between mice. The discrimination index (DI) was calculated as DI = (F_A_-F_x_)/(F_A_+F_x_), where F_A_ is the percentage of freezing in Context A and F_x_ is the percentage of freezing in either Context B or Context C.

### Nano-LC-MS/MS analysis

Cells were isolated as described in Section 2.2.4 and washed 3 times in 1X PBS prior to freezing in liquid nitrogen. Cell pellets were lysed in 10% sodium dodecyl sulfate (SDS) containing 100 mM triethylammonium bicarbonate (TEAB) with Pierce protease inhibitor (ThermoFisher Scientific) supplemented, sonicated, and centrifuged at 14,000 rpm for 10 minutes. Protein supernatant was collected and quantified using a BCA Protein Assay Kit (ThermoFisher). 100 ug of protein per sample was processed, trypsin digested in S-Trap microcolumns (Protifi), and peptides labeled according to the TMT10plex Isobaric Label Reagent Set (ThermoFisher) manufacture instructions. 100 fmol green fluorescent protein (GFP) per μg of protein was spiked in each sample before digestion. Each individual isobaric labeled sample was combined, lyophilized, and resuspended in 10 mM ammonium hydroxide prior to fractionation by high pH reversed-phase liquid chromatography. 60 fractions were collected and further concatenated into 20 fractions by combining 3 fractions for every 20 fractions apart(Yang et al., 2012). All fractions were dried down completely then resuspended in 0.1% formic acid. Peptide chromatographic separations and mass detection occurred with an Agilent 1260 nano/capillary HPLC system (Agilent Technologies) coupled to an Q-Exactive Orbitrap mass spectrometer (MS, ThermoFisher Scientific). Peptides were loaded onto an Acclaim PepMap 100 trap column (75 μm × 2 cm nanoViper, C18, 3 μm 100 Å, ThermoFisher Scientific) at flow rate of 2 μL/min. Peptides were further separated using a Zorbax 300SB-C18 column (0.075×150 mm, 3.5 μm 300 Å) (Agilent Technologies) at a flow rate of 0.25 μL/min, and eluted using a 5-30% mobile phase B gradient consisting of 0.1% formic acid in acetonitrile over 90 minutes. Data was collected in the data-dependent acquisition (DDA) mode at a mass resolution of 70,000 and scan range 375-2000 m/z. Automatic gain control (AGC) target was set at 1 × 10^6^ for a maximum injection time (IT) of 100 ms. The top 10 most abundant precursors (charge state between 2 and 5) were selected for MS/MS analysis. MS/MS spectra were acquired at a resolution of 35,000, AGC target at 1 × 10^6^, and maximum IT of 50 ms. All raw mass spectrometry data is publicly available on MassIVE with project ID MSV000090289 (ftp://massive.ucsd.edu/MSV000090289/).

The raw MS/MS data for each sample was searched against the curated SwissProt *Mus musculus* database in Proteome Discoverer software (v2.3.0.523, ThermoFisher Scientific), where trypsin was selected as the protease with 2 or less missed cleavage sites, precursor and fragment mass error tolerances set to 10 ppm and ± 0.02 Da, and only peptide precursors of +2, +3, +4 were analyzed. Peptide variable modifications allowed during the search were: oxidation ((+15.995 Da; M), TMT6 (+229.163 Da; S, T), and acetylation (+42.011 Da; N-terminus), whereas carbamidomethyl (+57.021 Da; C) and TMT (+229.163 Da; any N-terminus) were set as static modifications. Samples were grouped as iNSC Cav-1 KO (n=3) and iNSC Cav-1 WT (n=3). Protein identifications were accepted if they contained at least 2 unique peptides and abundances normalized to total peptide abundance. Differentially expressed proteins (DEP) for the iNSC Cav-1 KO NSPCs relative to the iNSC Cav-1 WT NSPCs were determined by applying ANOVA with a p-value of < 0.05.

### Protein pathway and mitochondria protein analysis

Protein pathway analysis was conducted on DEP using Ingenuity Pathway Analysis (IPA, Qiagen) and Gene Ontology (GO) enrichment analysis (Ashburner et al., 2000; Sandhu JK, 2021) using g:Profiler (Raudvere et al., 2019) (ve107_eg54_p17_bf42210). Functional enrichment maps were constructed in Cytoscape (Shannon et al., 2003) with EnrichmentMap pipeline collection applications (Reimand et al., 2019). Mitochondria pathways and functional groups were determined by manual curation using the MitoCarta 3.0 dataset (Rath et al., 2021).

### Mitochondria and membrane isolation

Cells were washed 3 times with 1X PBS and mitochondria were isolated according to the manufacturer instructions using the Mitochondria Isolation Kit (ThermoFisher). Briefly, cells were pelleted by centrifugation at 300g for 5 mins at 4°C. Reagent A containing protease and phosphatase inhibitor cocktails (ThermoFisher) was added to the cell pellet and placed on ice for 2 mins. Cells were then lysed by 100 strokes using a Dounce tissue homogenizer. Following the addition of Reagent C, the lysate was centrifuged at 700g for 10 mins at 4°C. The supernatant was then collected and centrifuged at 3000g for 15 min at 4°C to pellet mitochondria from the cytosol. The cytosol containing supernatant was collected and the mitochondria enriched pellet was washed with Reagent C and then centrifuged for 12000g for 5 mins at 4°C. The supernatant was centrifuged at 5.5 × 10^5^ rpm for 1 h to collect the membrane fraction. The pellets corresponding to mitochondria and plasma membrane were dissolved with 2% CHAPS in TBS and processed for BCA protein estimation and western blotting as described.

### TMRM Live Cell Imaging and mitochondria morphology analysis

Tetramethylrhodamine, methyl ester (TMRM) live cell imaging was conducted similarly as described(Jiang et al., 2022). Singly dissociated NPCs were grown on PLO/laminin coated glass bottom 24 well at a density of 50,000 cells/well. 5 days after plating, media was replaced with proliferation media containing 50 nM TMRM. Cells were incubated with TMRM for 30 mins at 37°C. Immediately prior to imaging, cells were washed 2 times with PBS followed by replacement of proliferation media lacking phenol red. 2D images were captured with a 63X objective using a Zeiss LSM880 META confocal microscope with heated stage and a DPSS 561-10 laser set at 0.1% laser power with GaAsP detection.

For quantification of mitochondria morphology, images were analyzed with ImageJ as described (Chaudhry et al., 2020). For optimal results, all images were pre- and postprocessed to reduce noise. The workflow and procedures for image processing and thresholding followed: 1) 2D images, operating on each slice in the stack, were preprocessed using “subtract background” (radiusL=L1 μm) to remove background noise; *2*) “sigma filter plus” (radiusL=L0.1 μm, 2.0 sigma) to reduce noise and smooth object signal while preserving edges; *3*) “enhance local contrast” (block sizeL=L64, slopeL=L2.0 for 2D) to enhance dim areas while minimizing noise amplification; and *4*) “gamma correction” (valueL=L0.80 for 2D) to correct any remaining dim areas. To identify mitochondria in the images, the “adaptive threshold” method was used. In the adaptive threshold plugin, block size was set to an equivalent of 1.25 μm. The thresholded images were then postprocessed using “despeckle” and then “remove outliers” (radiusL=L0.15 μm) to remove residual noise. Further to analyze mitochondrial 2D morphology, the image was first processed and thresholded (steps above), and the resulting binary image was used as the input for the “analyze particles” command (sizeL=L0.06 μm^2^-infinity, circularityL=L0.00–1.00), measuring for “area,” “perimeter,” and “shape descriptors.” For branch length analysis, the “skeletonize 2D/3D” command was applied to the thresholded image to produce a skeleton map, and the “analyze skeleton” command was used to calculate branch lengths in the skeletonized network.

For 2D time lapse videos, images were captured with 3x digital zoom for 120 s at frame rate of 3 Hz. Mitochondria velocity and total distance traveled were measured using the Trackmate (Ershov et al., 2022) plugin in ImageJ.

## Statistics

In all graphs, data is shown as mean ± SEM. Prism (Graphpad) was used for statistical analysis with tests indicated in figure legends. The following was used for p values: ns > 0.05, *p < 0.05, **p < 0.01, ***p < 0.001, and ****p < 0.0001.

## Acknowledgments

We thank Dr. Trongha Phan for his expertise and guidance on behavioral experiments and scientific discussion. We thank Dr. Pavan Kumar, Dr. Muskan Gupta, Abhi Ramakrishnan, Stephanie Dunning for help with animal husbandry, genotyping, and IP injections. We thank Karen Rakowiecki for assistance with cell culture experiments. We thank Dr. Christian Peters and Shana Netherton for use of Elevated Plus Maze equipment. We thank Dr. Ying Jiang for expertise and guidance on the mitochondrial assays. We also thank Dr. Peter Toth and Dr. Ke Ma at the UIC Fluorescence Imaging Core for guidance on microscopy imaging, Figen Seiler at the UIC Electron Microscopy Core for TEM imaging, and Dr. Balaji Ganesh at the UIC Flow Cytometry Core for assistance with the Mitochondria Seahorse Assay. Graphical abstract, Figure 2A, 3A, and 4E were created Biorender.com. This work was financially supported by National Institutes of Health, National Institution on Aging (NIA) F30AG071144 (TKLS), T32AG067468, AG033570, AG076940, AG062251, AG060238 (OL), NINDS/NIA:R01NS114413 (SMC) and Together Strong NPC Foundation (SMC).

## Author Contributions

TKLS and DP contributed equally to the design and execution of experiments, analysis of the data, writing and revisions of the manuscript. LA, EQ, YI, KO, EQ, PK, JM and WL assisted with data collection and analysis. AS and JB assisted with the design of the studies and generation of the mouse models used. RDM and SMC assisted with the data analysis, interpretation, and revision of the manuscript. OL supervised the project, assisted with the data analysis, interpretation and revised the manuscript. All authors contributed to the review of the manuscript.

---

**Figure S1.** Generation of inducible Nestin specific Cav-1 knockout (iNSC Cav-1 KO) mouse model. iNSC Cav-1 KO mice displays altered levels of NSC proliferation yet no changes immature neuron formation at 3 months of age. (A) Schematic of NestinCre^ERT2/+^;Cav-1^fl/fl^ mice, where mice were injected at 4-5 weeks of age with either corn oil (Corn) or tamoxifen (TAM) for 5 consecutive days to generate control mice (iNSC Cav-1 WT) and Cav-1 knockout mice (iNSC Cav-1 KO), respectively. (B) RT-qPCR quantification of Cav-1 transcript expression from NSPC isolated from the iNSC Cav-1 KO and iNSC Cav-1 WT mice. (C) Cav-1 immunoblot, and quantification normalized to β-actin of protein lysate from NSPC isolated from the iNSC Cav-1 KO and iNSC Cav-1 WT mice. (D) Immunocytochemistry of Cav-1 and confocal imaging of Cav-1 (green) and DAPI (blue) in hippocampal NSPC isolated from iNSC Cav-1 WT and iNSC Cav-1 KO mice shown. Scale bar, 20 μm. (E) Ultrastructural electron micrographs of caveolae and clathrin coated vesicles in hippocampal NSPC isolated from iNSC Cav-1 WT and iNSC Cav-1 KO mice. Solid green arrowhead indicates caveolae. Open green arrowhead indicates clathrin coated vesicle. Scale bar, 200 nm. (F) Representative images of TBR2 (red), ASCL1 (green), and DAPI (blue) markers in the DG of iNSC cav-1 WT and iNSC Cav-1 KO mice at 3 months of age. Yellow arrowheads indicate TBR2^+^ASCL1^+^ cells. Scale bar, 50 µm. (G-I) Quantification of total intermediate progenitor cells (TBR2^+^ ASCL1^+^) in the DG of iNSC Cav-1 WT and iNSC Cav-1 KO mice at 3 months of age. n=4 mice per group. (J) Representative images of TBR2 (red), ASCL1 (green), and DAPI (blue) positive cells in the DG of iNSC cav-1 WT and iNSC Cav-1 KO mice at 6 months of age. Yellow arrowheads indicate TBR2^+^ASCL1^+^ cells. Scale bar, 50 µm. (K-M) Quantification of total intermediate progenitor cells (TBR2^+^ ASCL1^+^) in the DG of iNSC Cav-1 WT and iNSC Cav-1 KO mice at 6 months of age. n=4 mice per group. (N-O) EdU uptake assay in NSPC isolated from iNSC Cav-1 WT and iNSC Cav-1 KO mice. Quantification of the percentage of EdU+ cells to the total DAPI between NSPC isolated from iNSC Cav-1 WT and iNSC Cav-1 KO mice. Scale bar, 20 μm. (P) Representative confocal images of NeuN (red), DCX (green) and DAPI (blue) in the DG of iNSC Cav-1 WT and iNSC Cav-1 KO mice at 3 months of age. Scale bar, 25 μm. (Q-S) Quantification of total DCX expressing cells, NPC and Neuroblasts (DCX^+^NeuN^-^) and immature neurons (DCX^+^NeuN^+^) in the DG of iNSC Cav-1 WT and iNSC Cav-1 KO mice at 3 months of age. n=5-6 mice per group. Data represented as mean ± SEM. Data analyzed by unpaired two-tailed Student’s t-test. ns p> 0.05, *p< 0.05, **p < 0.01, ***p < 0.001 ****p < 0.0001.

**Figure S2.** iNSC Cav-1 KO mice display no anxiety-like behavior or altered performance in a context generalization or spatial recognition learning and memory task compared to iNSC Cav-1 WT mice at 6 months of age. (A) Schematic of contextual fear generalization paradigm between Context A and C. See Materials and methods for details. (B-C) Quantification of percent freeze (30 min post-shock) and discrimination index in context A and C on Day 2 between the NSC Cav-1 WT and iNSC Cav-1 KO mice. n=12 iNSC Cav-1 WT and n=8 iNSC Cav-1 KO. (D-E) Quantification of percent freeze (24 hr post-shock) and discrimination index in context A and C on Day 3 between the NSC Cav-1 WT and iNSC Cav-1 KO mice. n=12 iNSC Cav-1 WT and n=8 iNSC Cav-1 KO. (F) Schematic of elevated plus maze (EPM). Mice were placed in the center with head towards the open arm and allowed to explore for 5 mins. See Materials and methods for details. (G-I) Quantification of time spent in the open arm, number of entries in the open arm and total distance traveled in the EPM of iNSC Cav-1 WT and iNSC Cav-1 KO mice. n=7 per genotype. (J) Schematic of novel object location (NOL). Mice were trained during two 10 min trials on day 1 followed by a 5 min testing trial on day 2 with one object moved to a new location. See Materials and methods for details. (K-L) Quantification of the percentage of time spent with objects in each location and the discrimination index determined between new and old locations on Day 2 of task between the iNSC Cav-1 KO and iNSC Cav-1 WT mice. n=7 per iNSC Cav-1 WT and n=5 iNSC Cav-1 KO. Data represented as mean ± SEM. Data analyzed by unpaired two-tailed Student’s t-test. ns p > 0.05, **p < 0.01, ***p < 0.001.

**Figure S3.** Loss of Cav-1 in mouse hippocampal NSPS alters trafficking dynamics. (A) Representative live-cell images of TMRM staining in NSPCs isolated from iNSC Cav-1 KO and iNSC Cav-1 WT mice. NSPCs were incubated with 50 nM TMRM for 30 min followed by confocal microcopy visualization. Mitochondria were skeletonized in ImageJ. Scale bar, 10 μm. (B-D) Quantification of mitochondria area per cell, mitochondria perimeter per cell, and circularity per cell in iNSC Cav-1 WT and iNSC Cav-1 KO NSCPs. N=25 cells per group. (E) Quantification of skeletonized mitochondria branch length per mitochondria in iNSC Cav-1 WT and iNSC Cav-1 KO NSCPs. N=25 cells per group. (F) Quantification of TMRM fluorescence in iNSC Cav-1 WT and iNSC Cav-1 KO NSPC. N=25 cells per group. (G) Representative images of mitochondria trafficking per minute using Trackmate (Ershov et al., 2022) in iNSC Cav-1 WT and iNSC Cav-1 KO NSPCs. Scale bar 10 μm. (H) Quantification of mitochondria velocity. n=10 cells per genotype with n=948 particles (mitochondria) for iNSC Cav-1 WT and n=1259 particles (mitochondria) for iNSC Cav-1 KO analyzed. (I) Quantification of mitochondria total distanced traveled. n=10 cells per genotype with n=948 particles (mitochondria) for iNSC Cav-1 WT and n=1259 particles (mitochondria) for iNSC Cav-1 KO analyzed. Data represented as mean ± SEM. Data analyzed in by unpaired two-tailed Student’s t-test. ns p > 0.05, *p < 0.05, **p < 0.01, ***p < 0.001 and ****p < 0.0001.

**Table S1.**
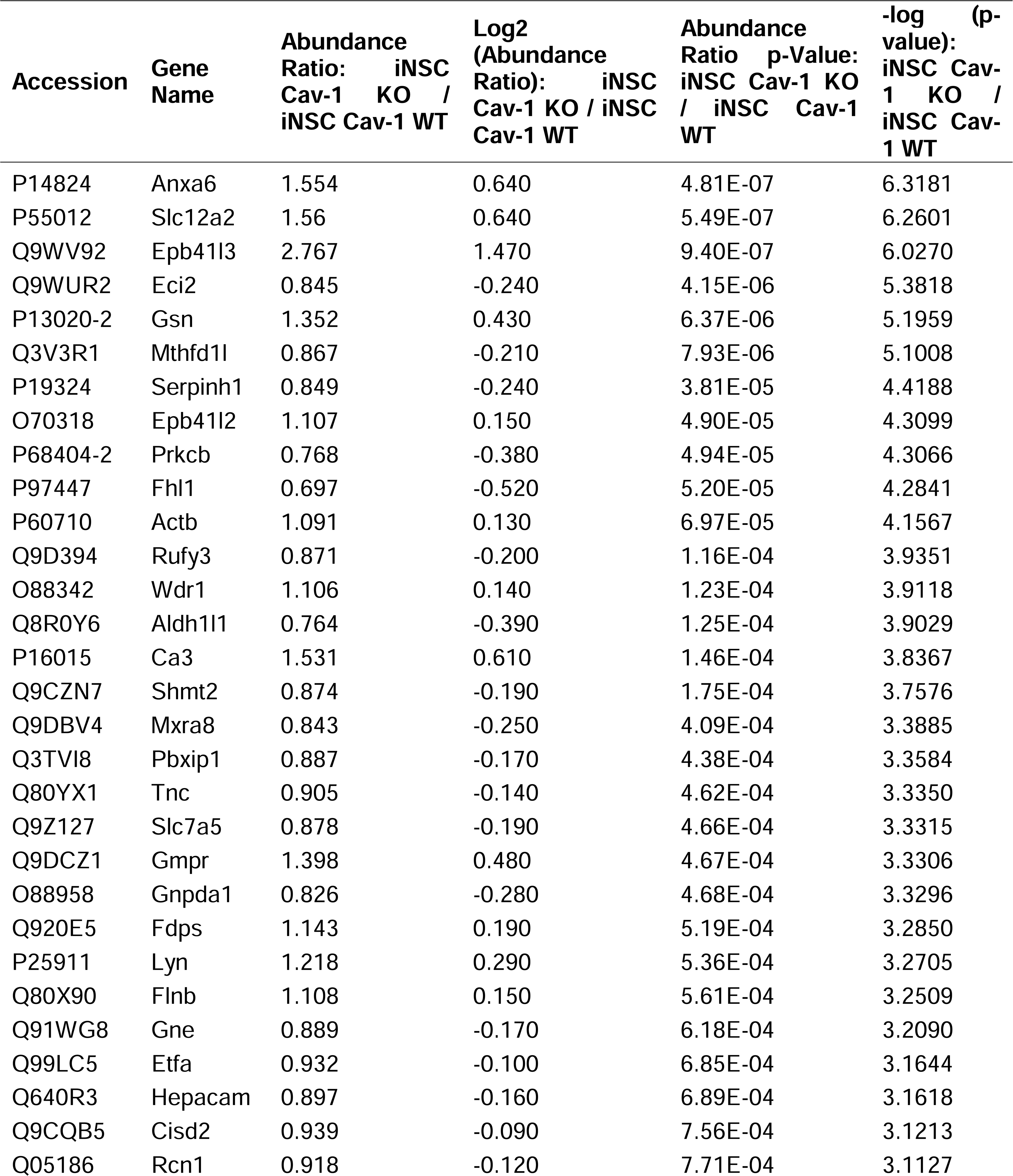
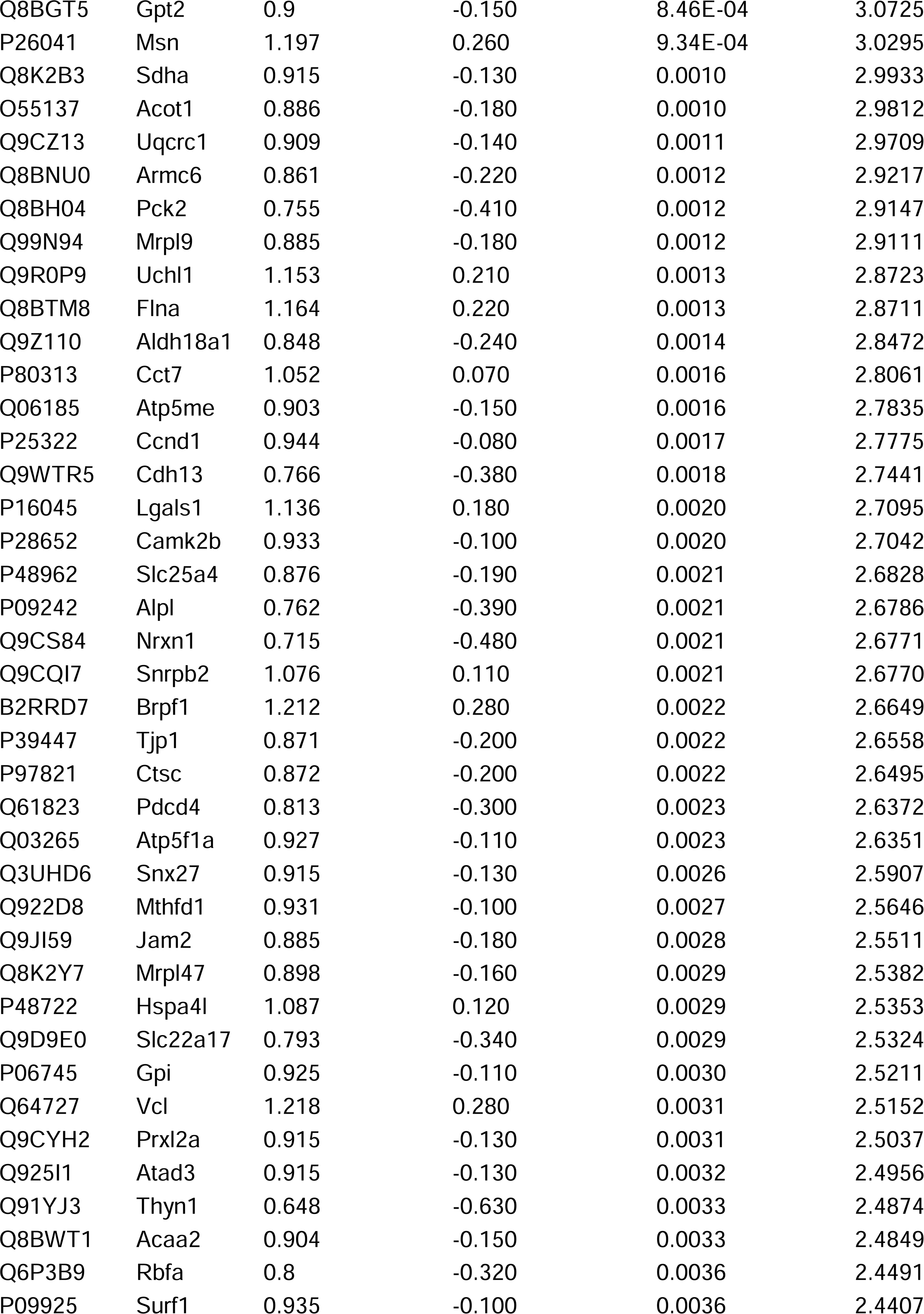
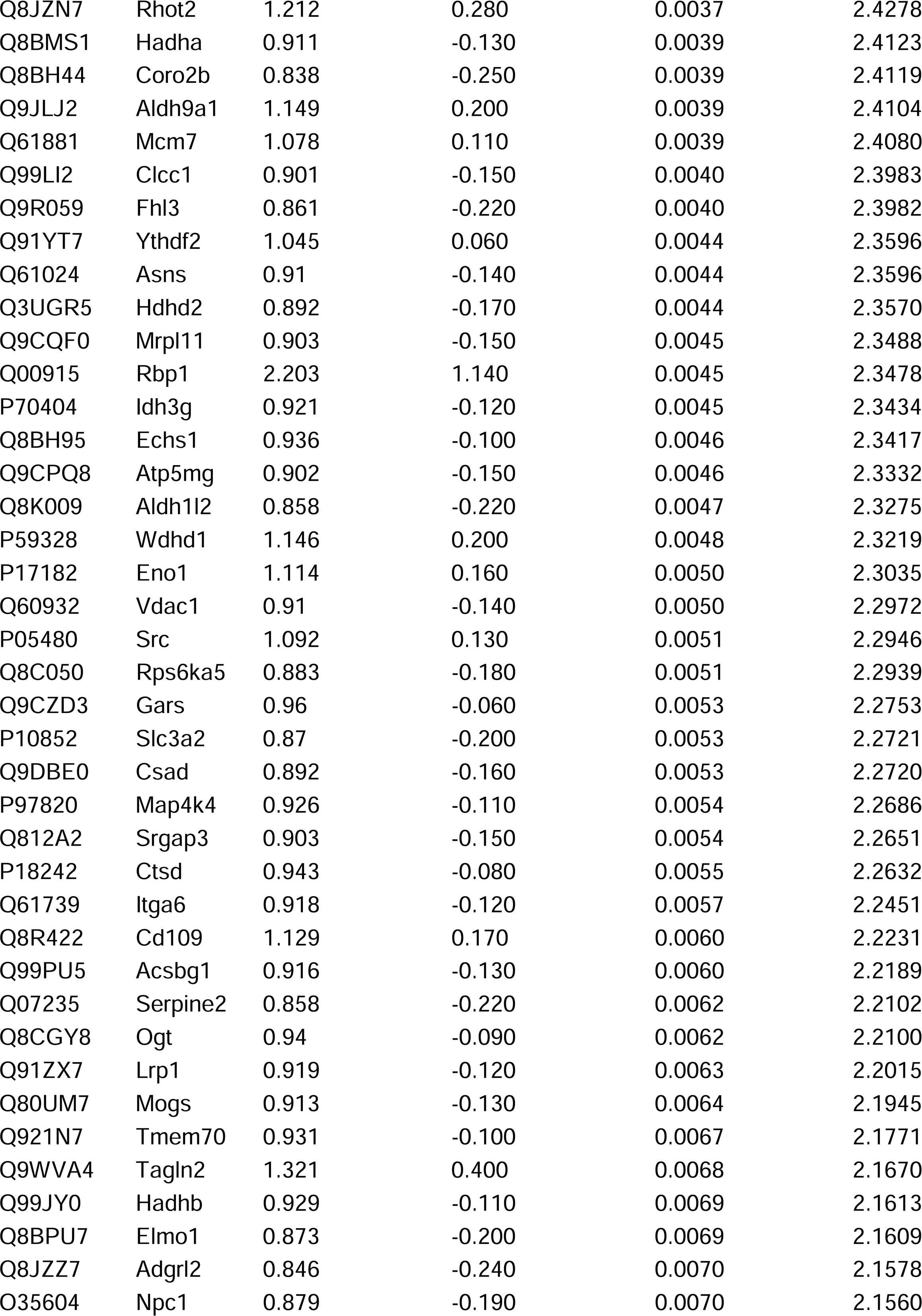
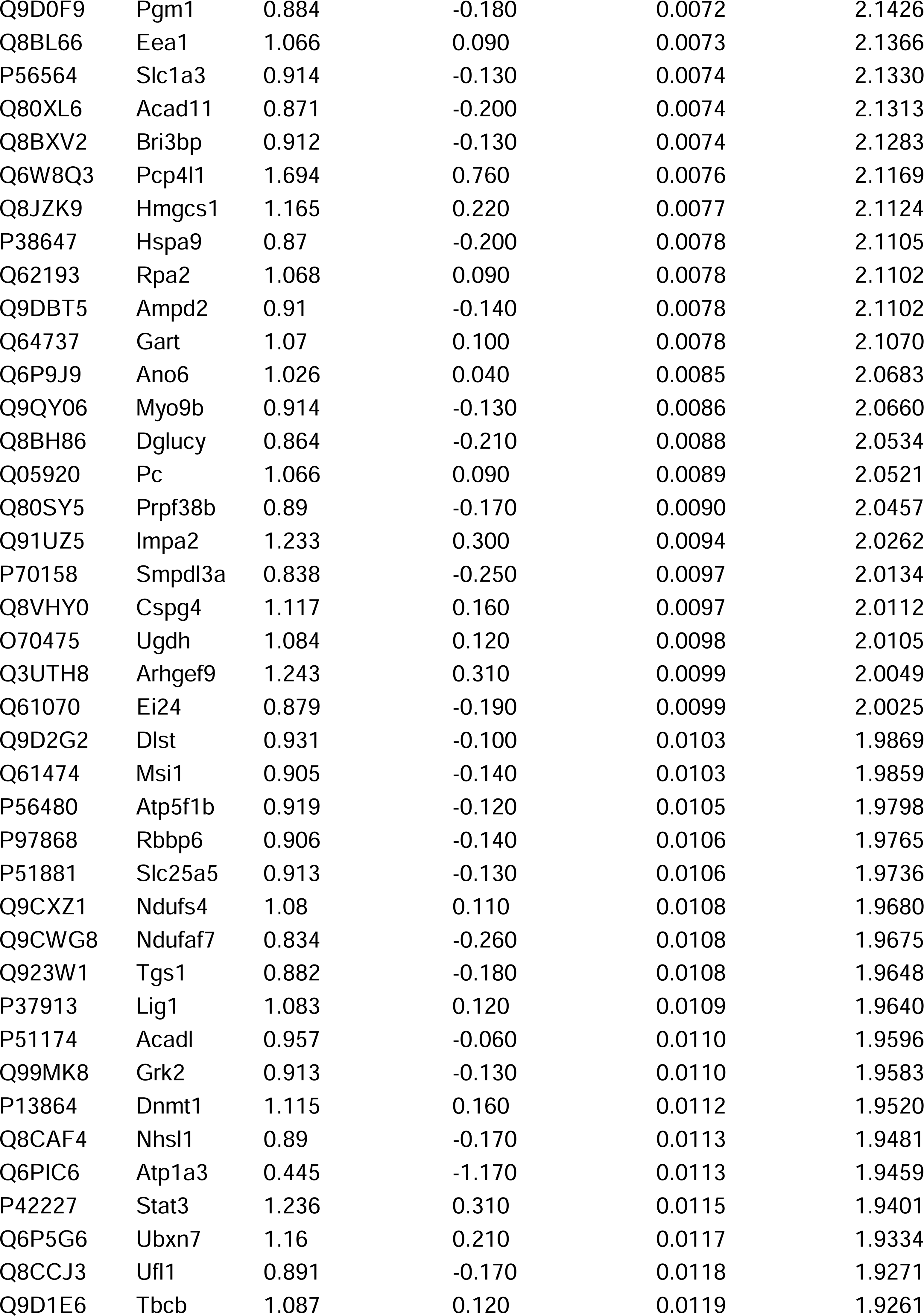
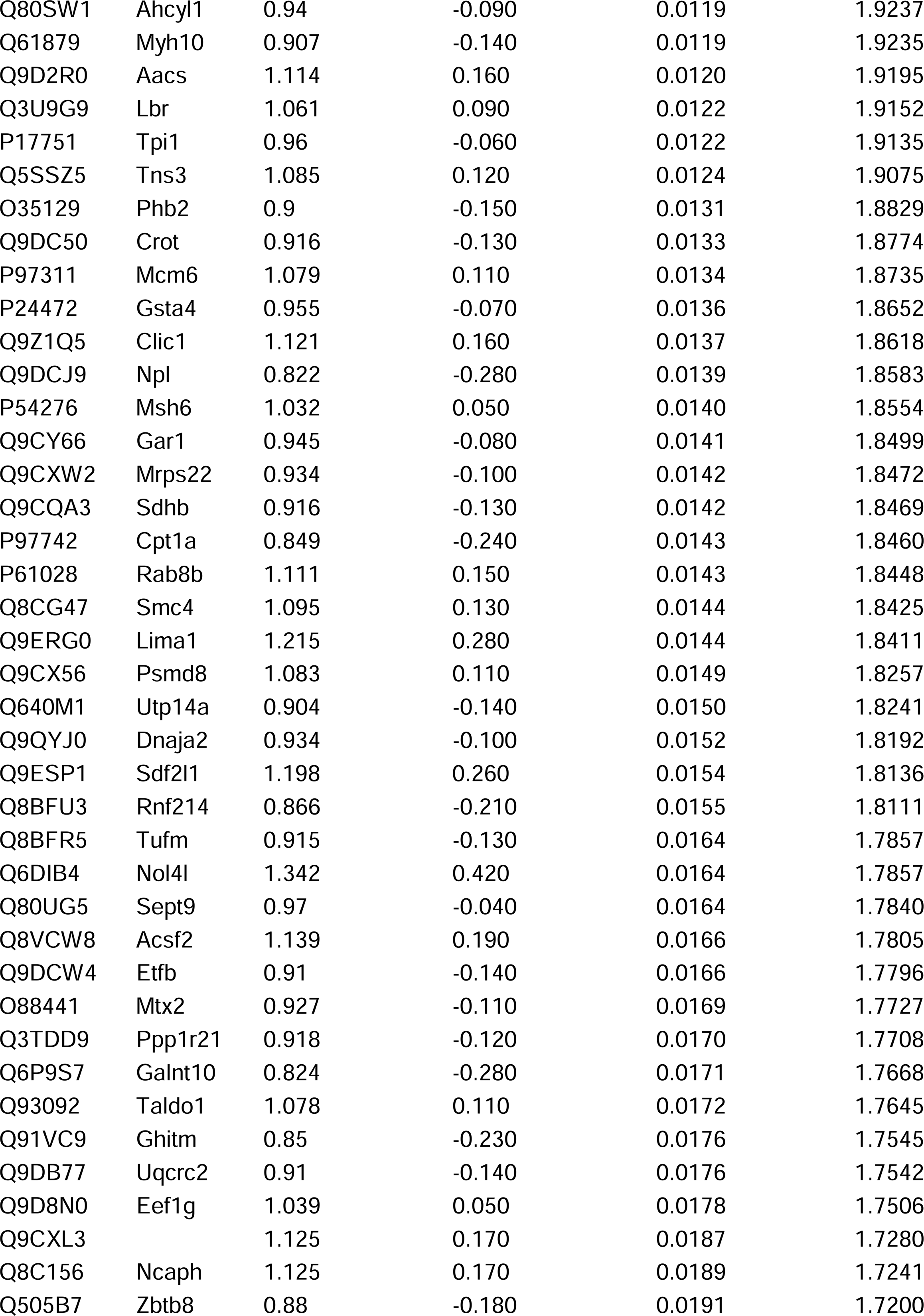
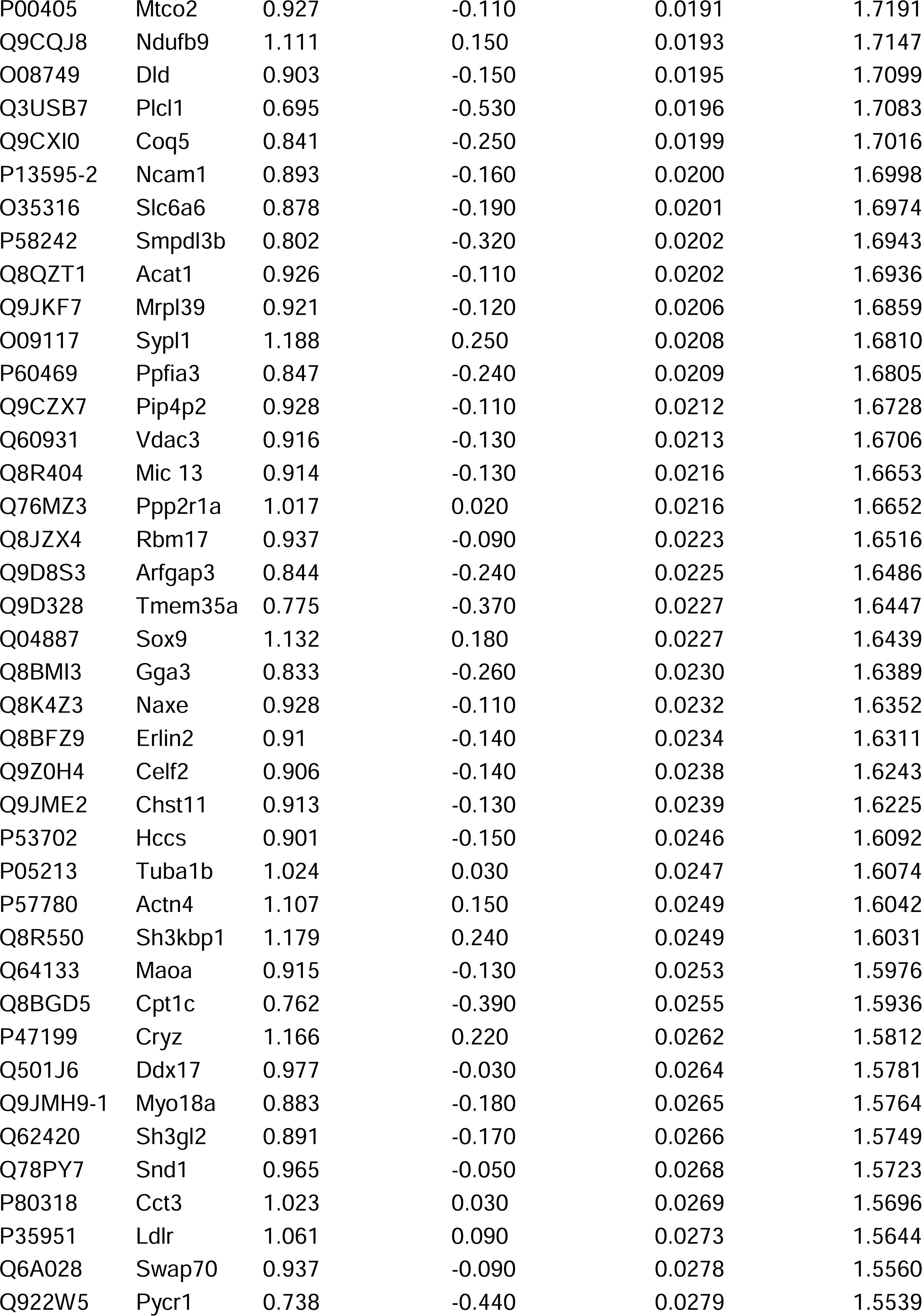
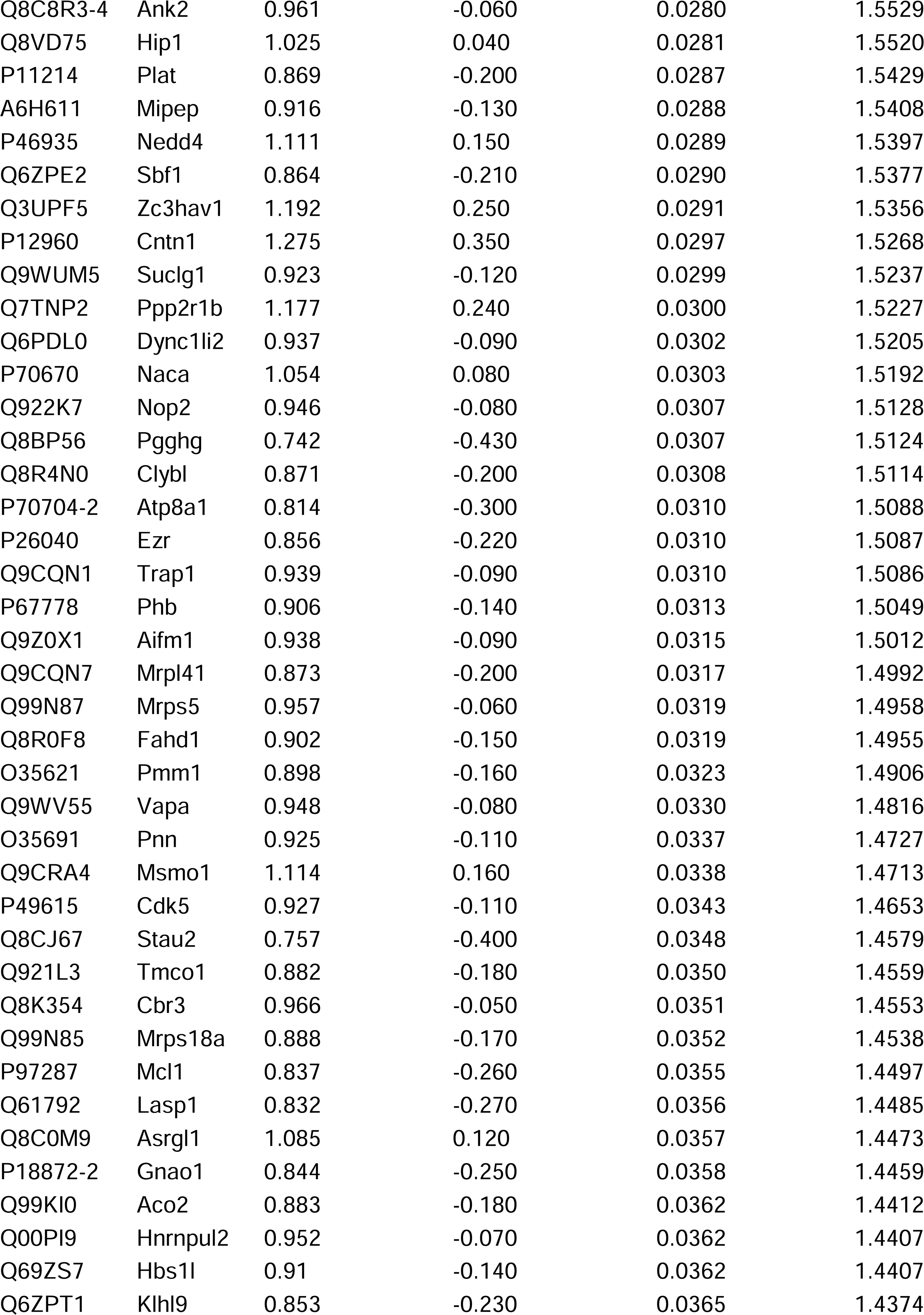
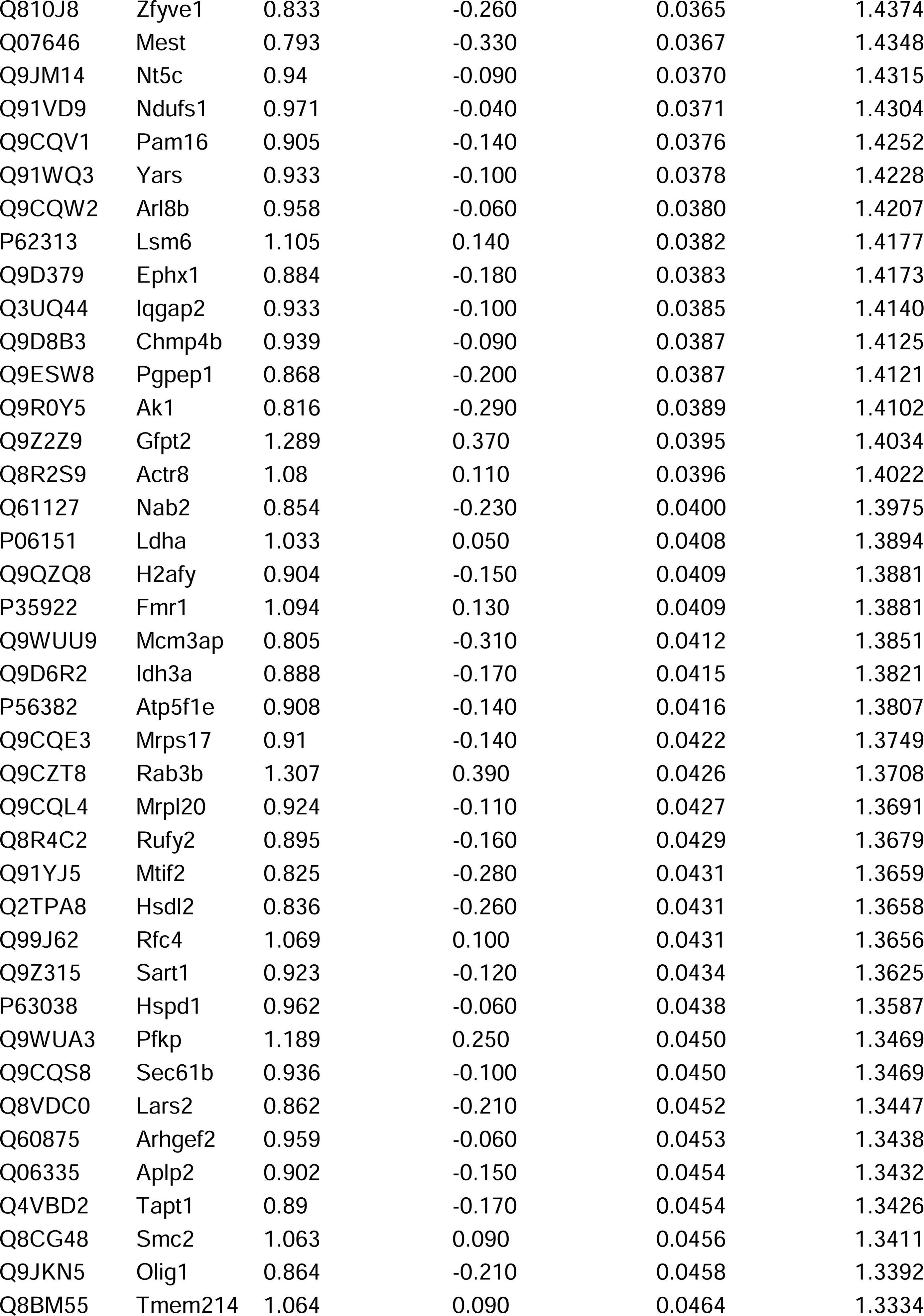
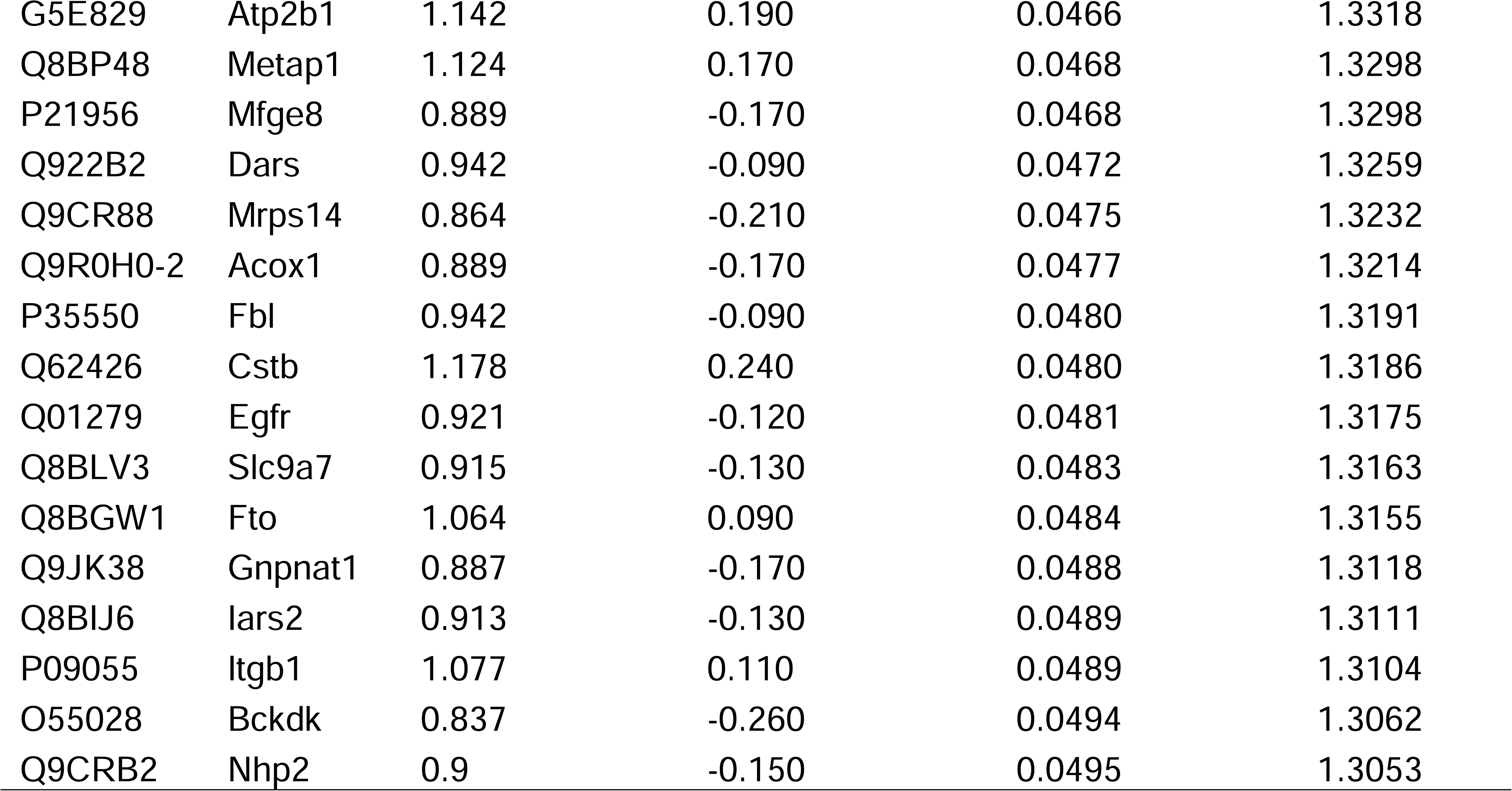
Differentially Expressed Proteins in iNSC Cav-1 KO vs iNSC Cav-1 WT hippocampal NSPCs (ANOVA p<0.05).

**Table S2.**
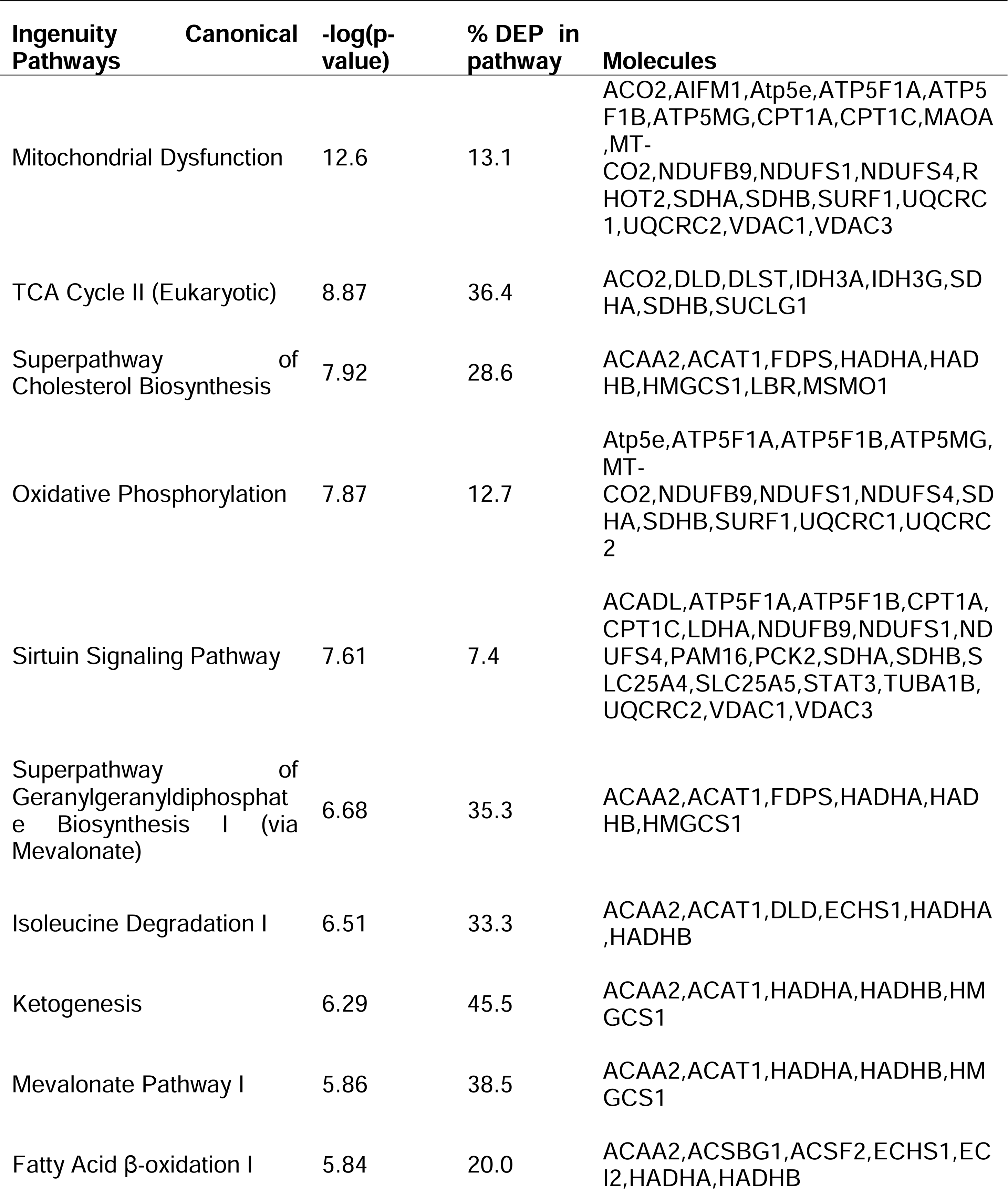
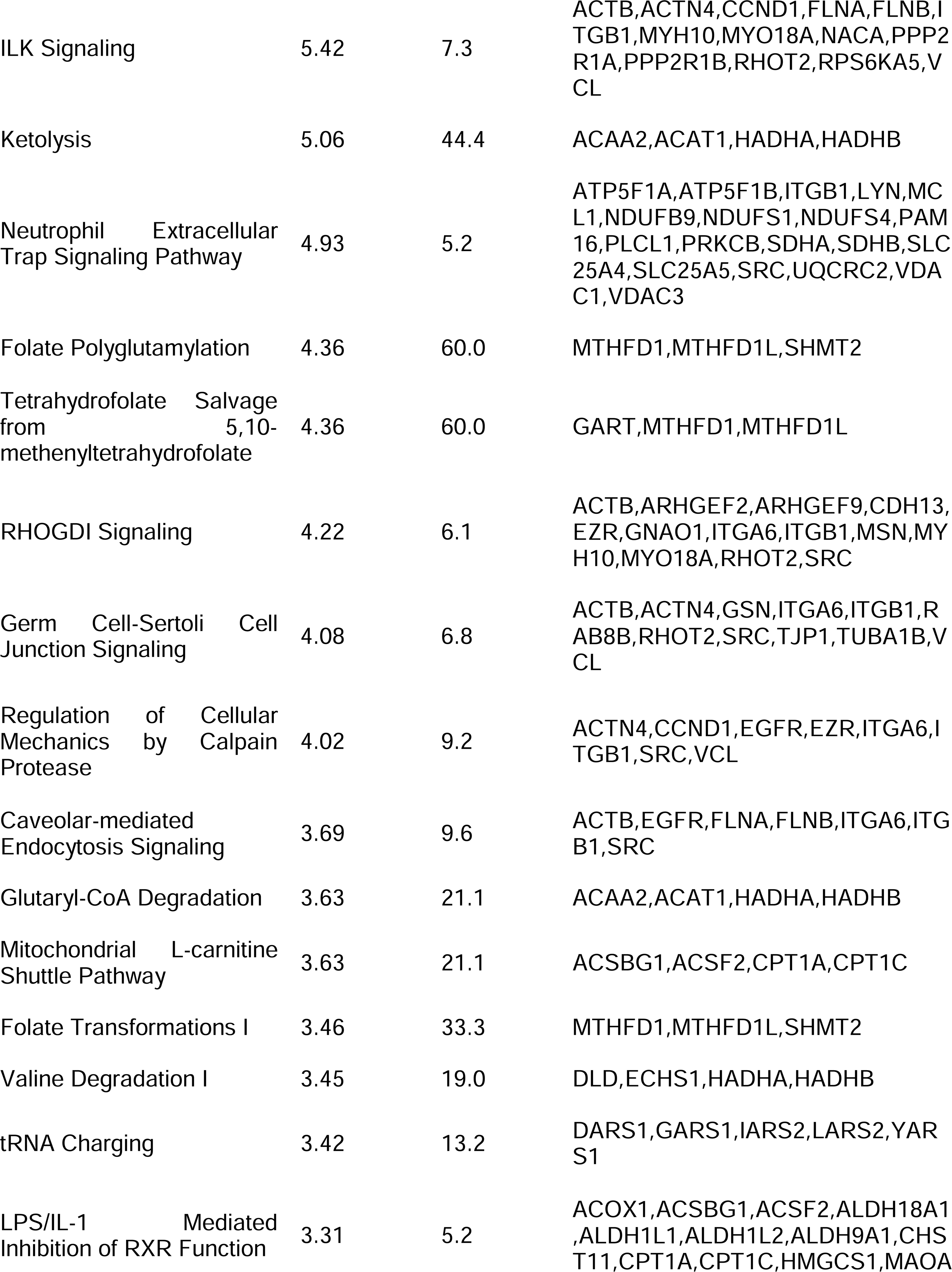
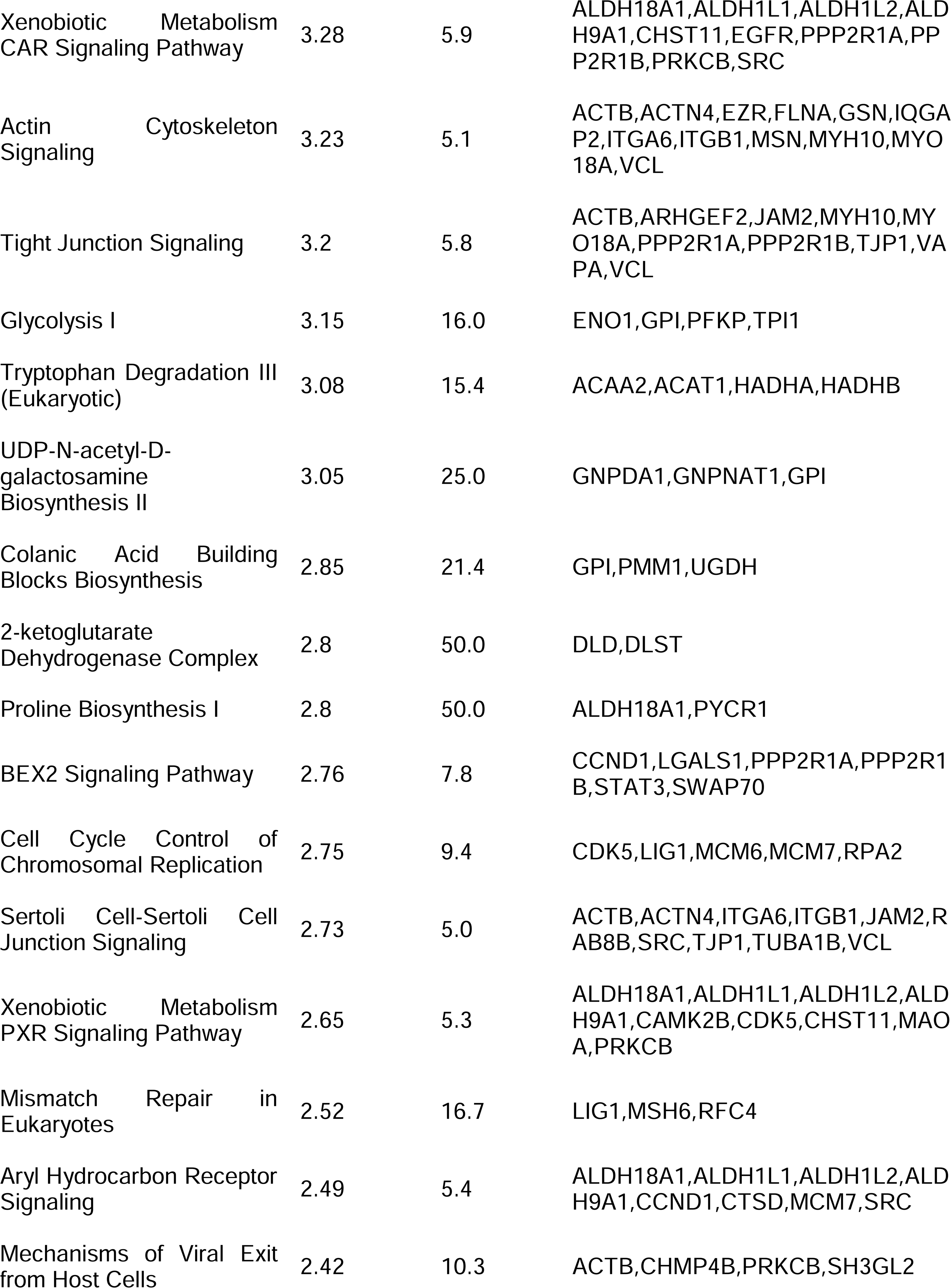
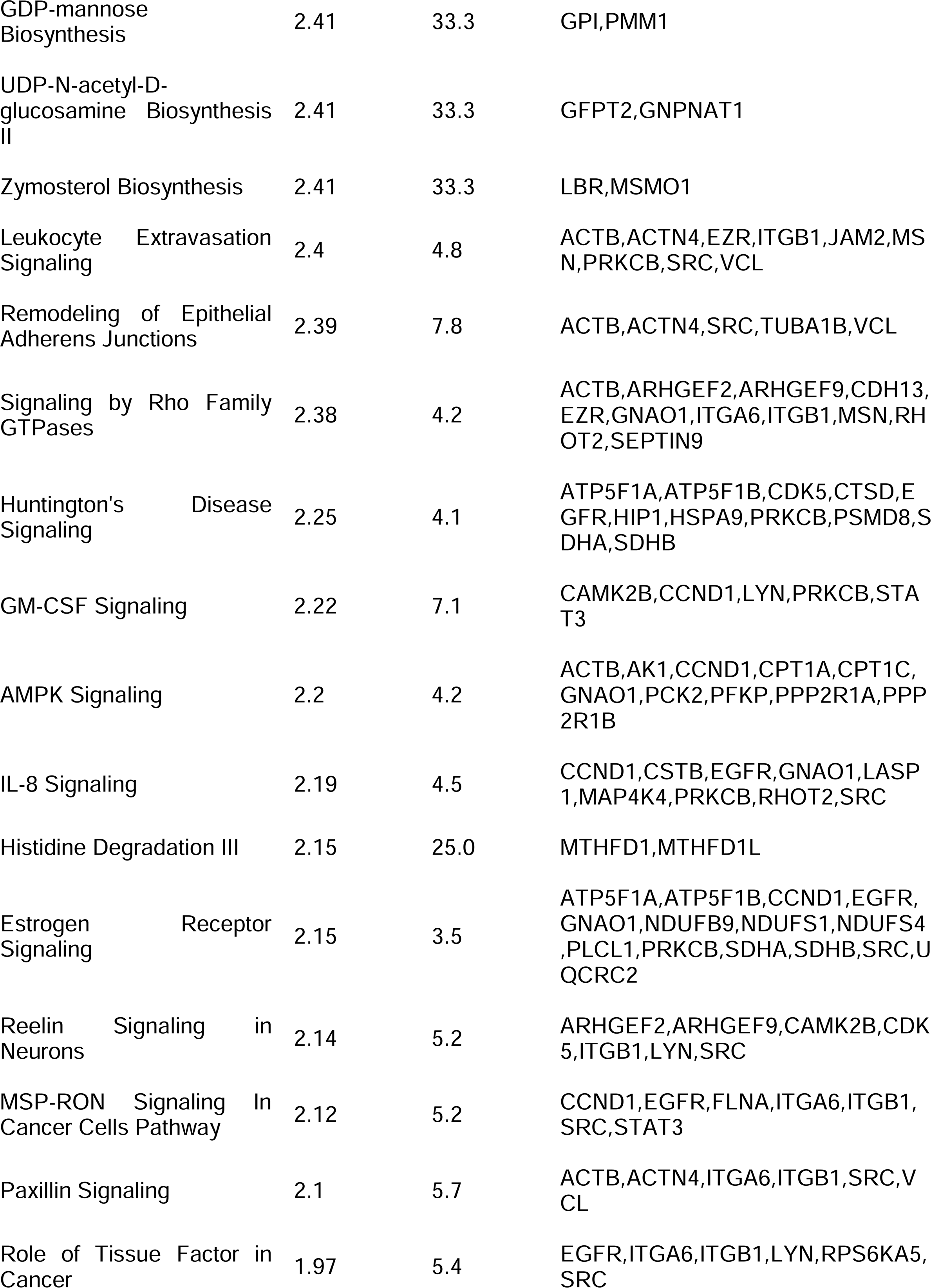
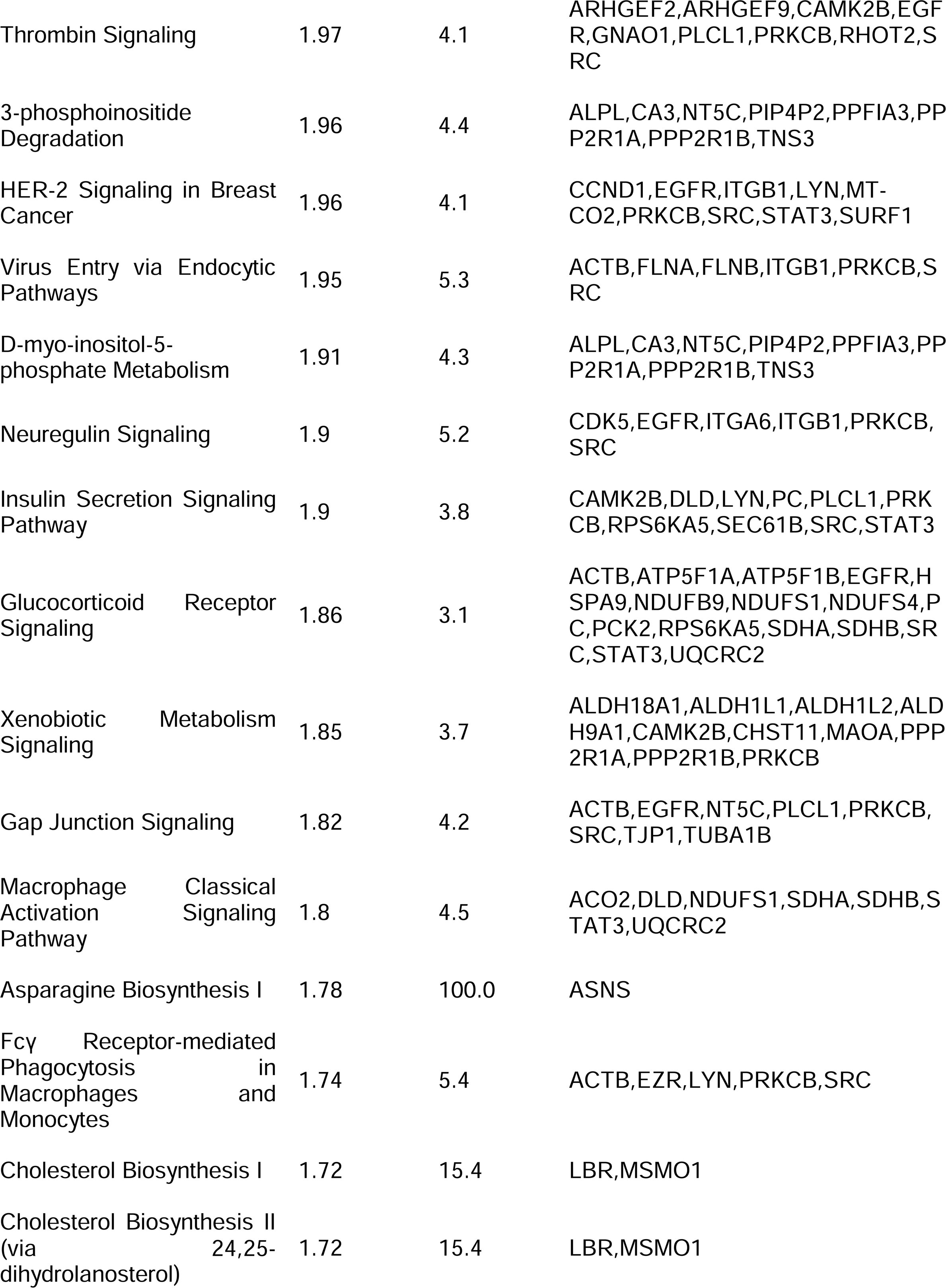
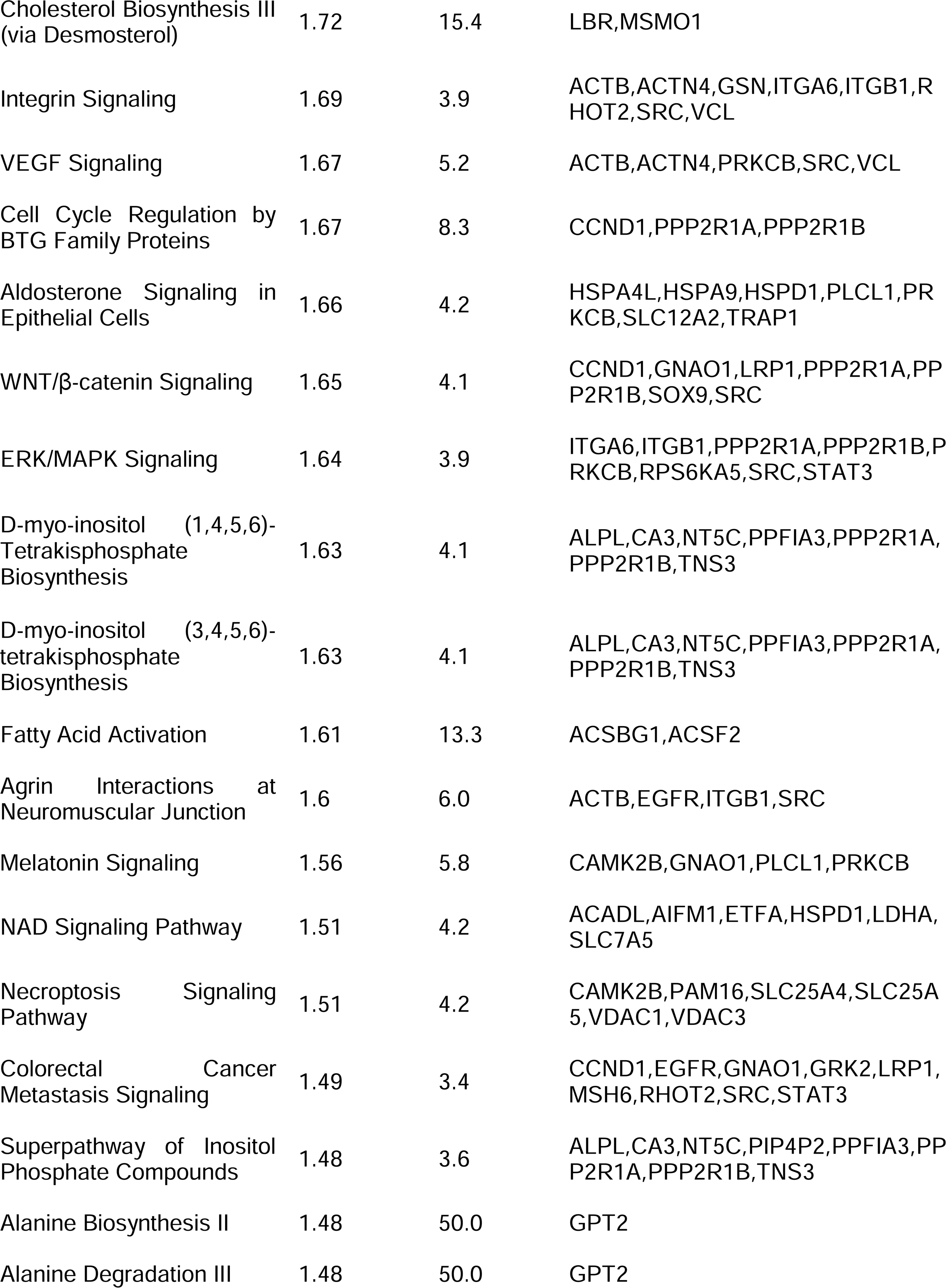
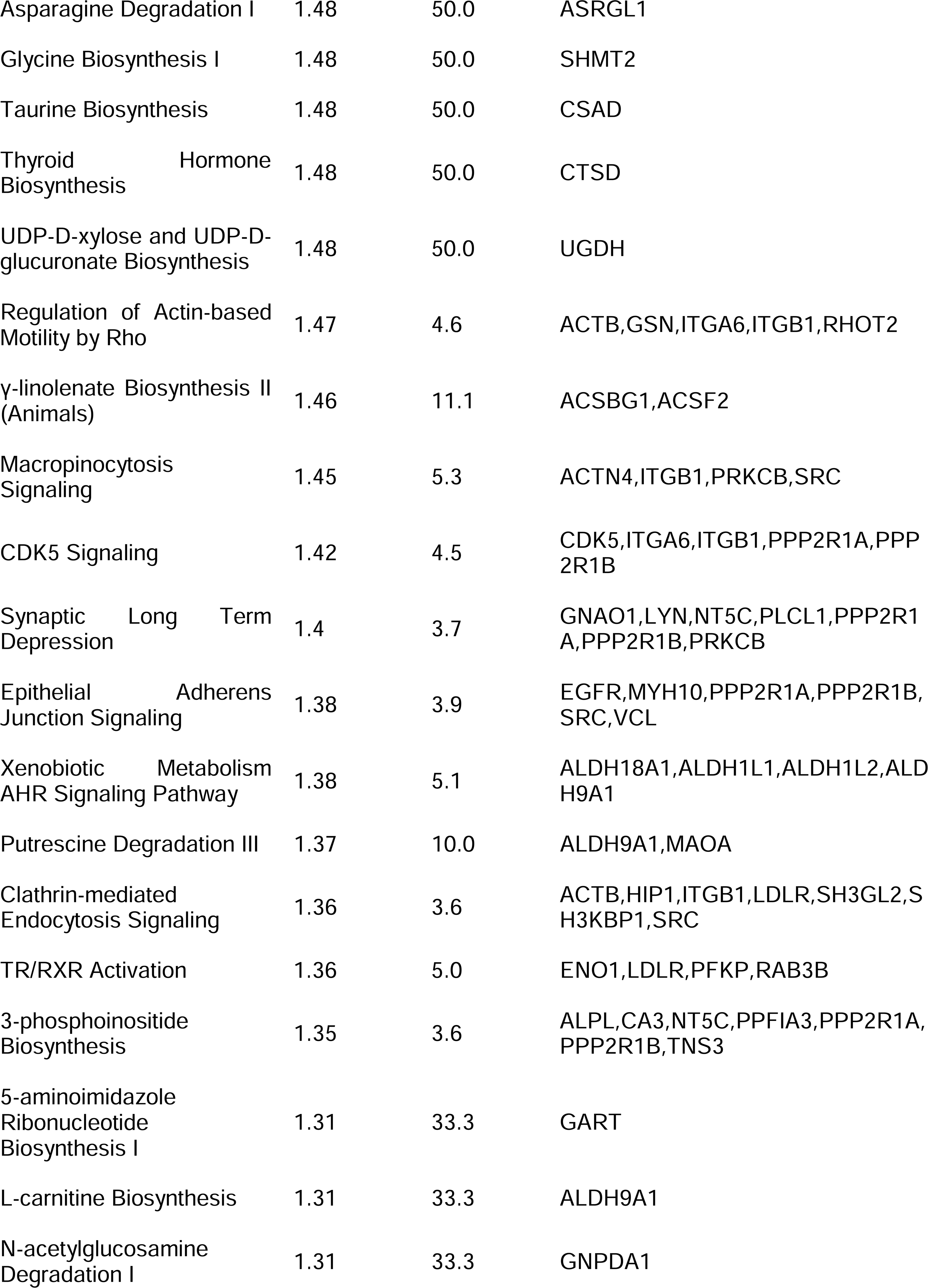
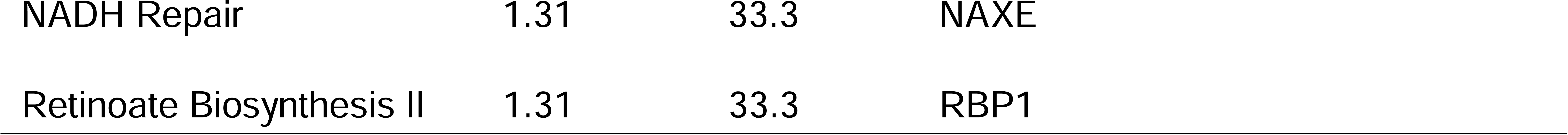
Ingenuity Pathway Analysis (IPA) of differentially expressed proteins (DEP) in iNSC Cav-1 KO vs Cav-1 WT hippocampal NSPCs.

