## Supplementary figures and images for "Caveolin-1 Regulates Neurogenesis and Learning and Memory by Modulation of Mitochondrial Dynamics"

### Supplemental Figure 1

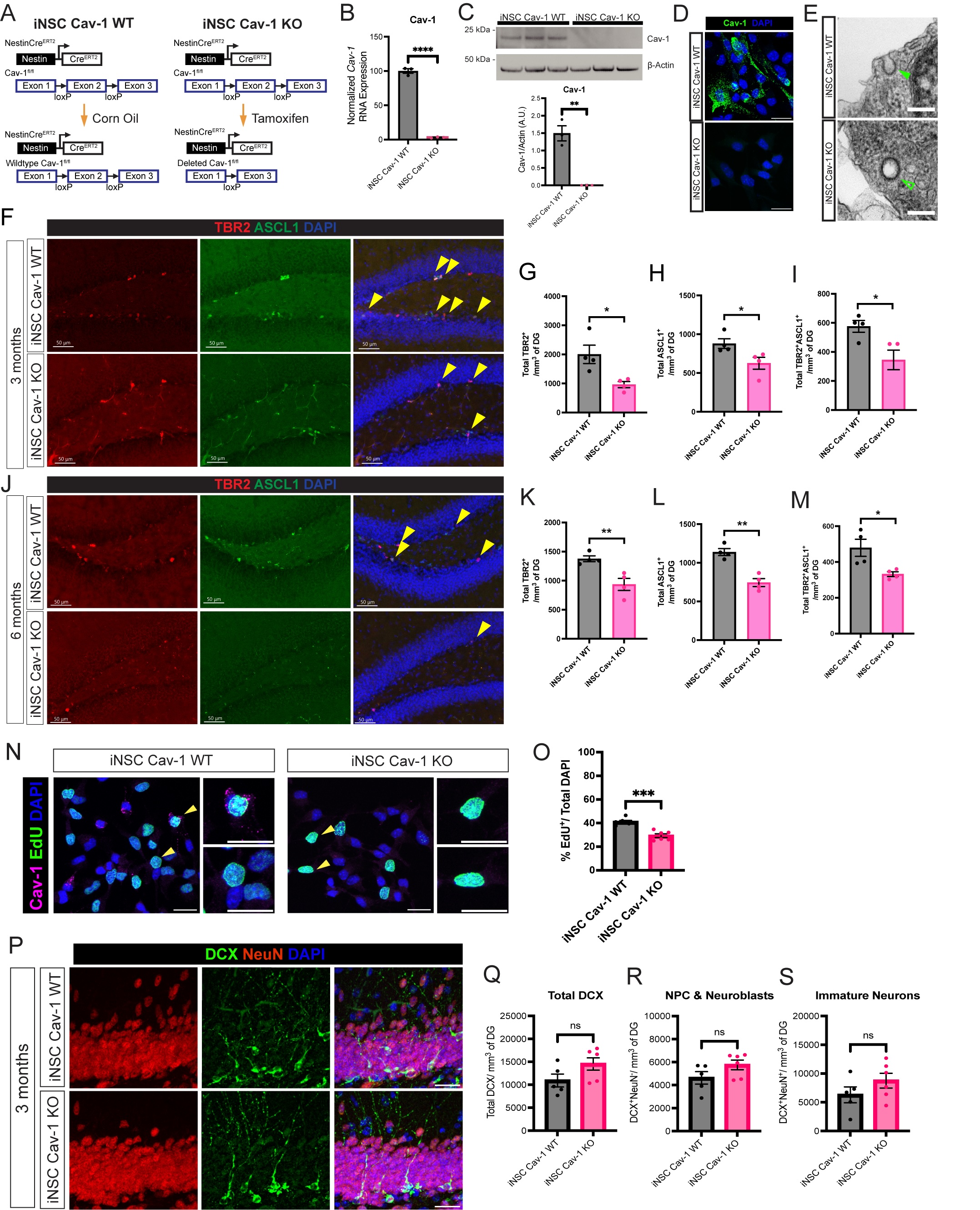

### Supplemental Figure 2

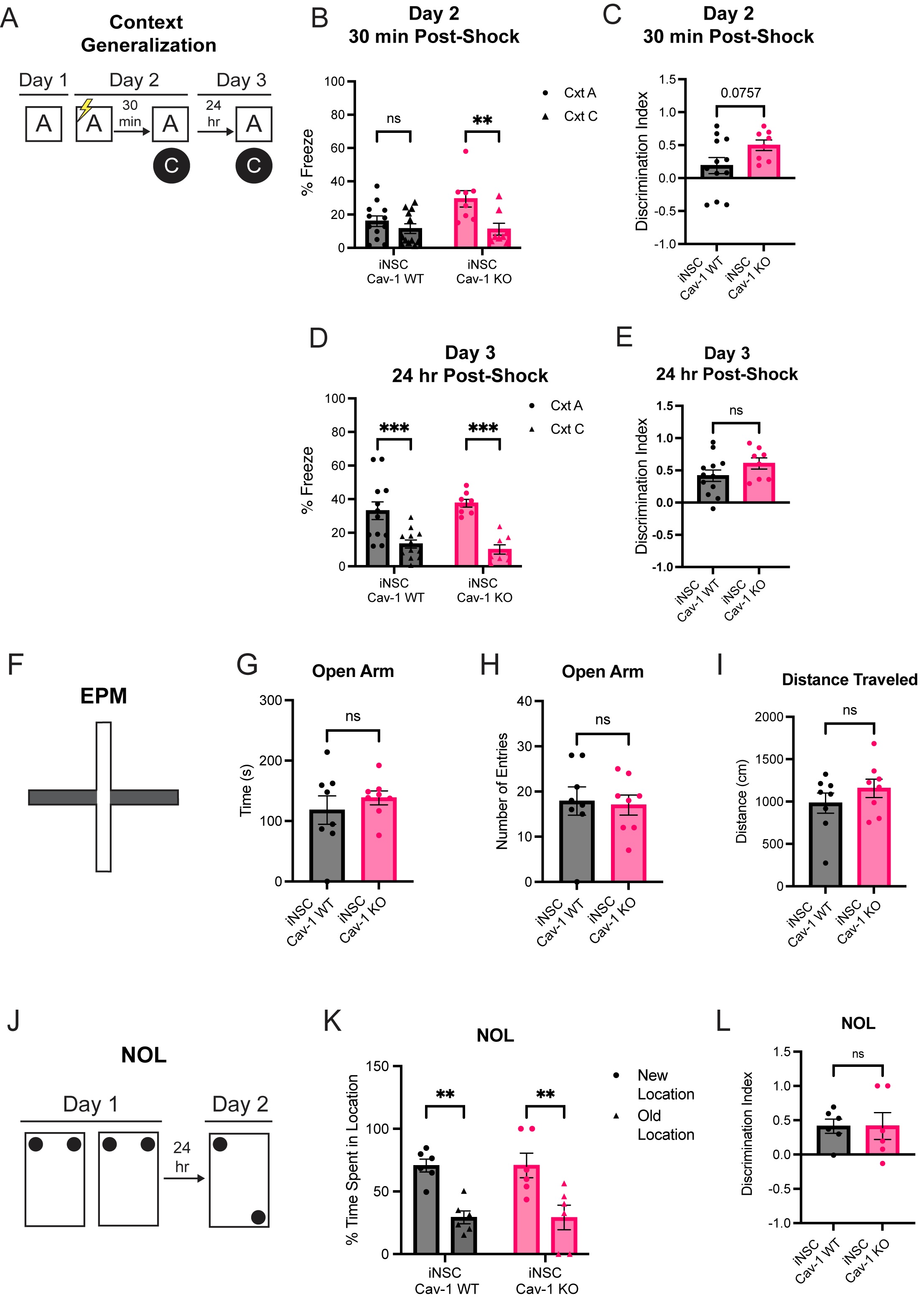

### Supplemental Figure 3

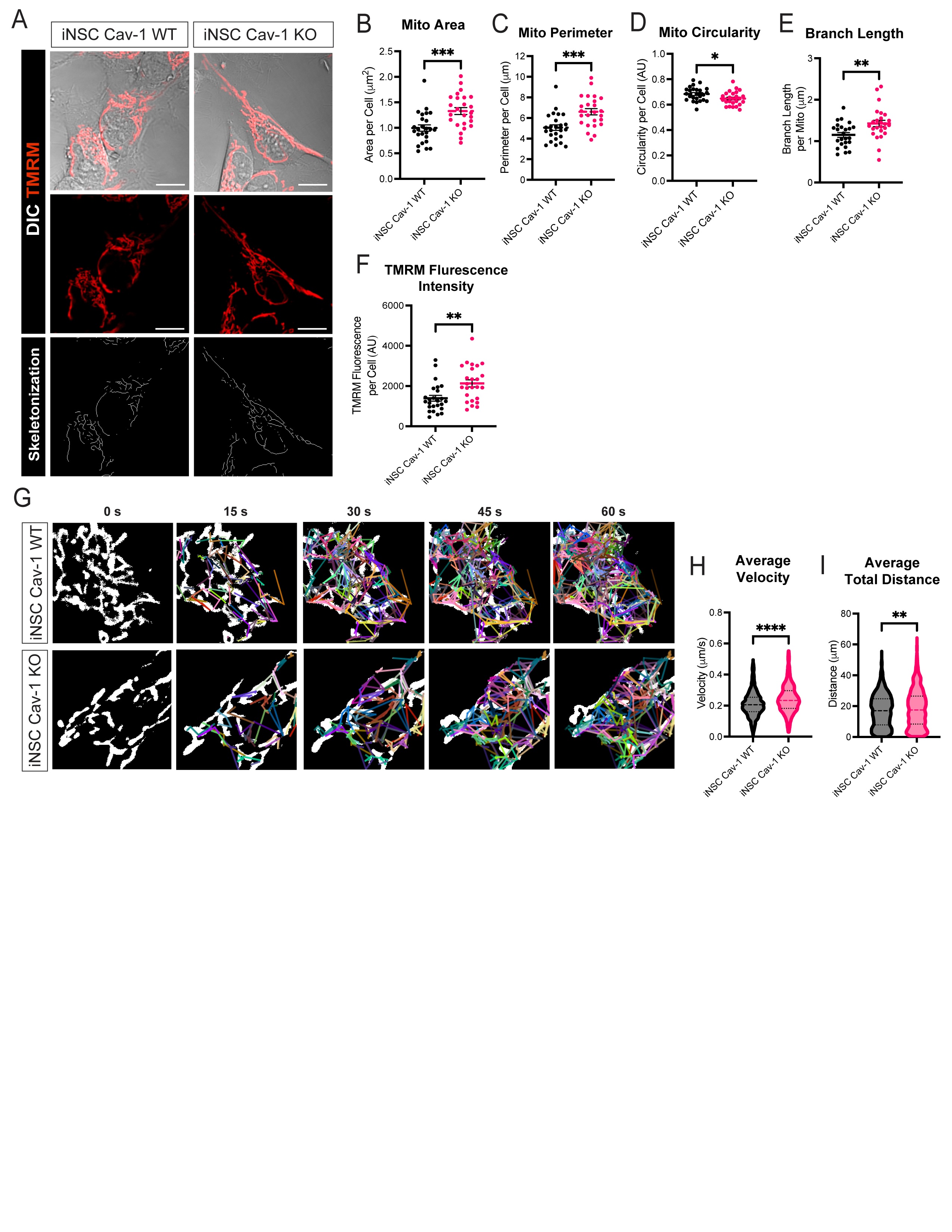
